# High-level visual cortex representations are organized along visual rather than abstract principles

**DOI:** 10.1101/2024.11.12.623145

**Authors:** Adva Shoham, Rotem Broday-Dvir, Rafael Malach, Galit Yovel

**Affiliations:** School of Psychological Sciences, Tel Aviv University; Department of Brain Sciences, Weizmann Institute of Science; Sagol School of Neuroscience, Tel Aviv University

## Abstract

A fundamental question dominating the study of human visual cortex is whether it is organized along visual or semantic information. This question is unresolved, and the controversy has been rekindled by the recent report that surprisingly, revealed that the representations of textual description of images by linguistic artificial networks’ successfully predict the response of high-level visual cortex to visual images. These findings appear to support a linguistic, abstract, organizing principle of human visual cortex. Here, using iEEG recordings from high level visual cortex in patients, we contributed to this debate, by testing the hypothesis that this linguistic alignment is restricted to textual descriptions of the visual content of the images (visual text) and does not extend to abstract textual descriptions (abstract text). We selected images that depict familiar faces and places, as these images allow for the best dissociation between these two types of text and generated their visual and abstract (e.g., name and biography of a person) textual descriptions. We then predicted the relational structures of the iEEG response to the images using their textual representations based on a large language model and the image representation based on a convolutional neural network. Neural relational-structures in high-level visual cortex were similarly predicted by images and visual-text but not abstract-text representations. Abstract text best predicted responses of the fronto-parietal cortex to the images. These results demonstrate that visual-language alignment in high-level visual cortex is limited to visually grounded language.

## Introduction

High-level visual cortex shows selective responses to different categories, including objects, scenes, faces, and bodies^1–7^. These category selective responses are localized in distinct cortical regions, with animate stimuli, such as faces and bodies, more laterally, and inanimate objects and scenes more medially (for review see^3^, but see^8,9^ for a different perspective). While there is substantial consensus with regards to the anatomical locations of the various visual representations-the principles by which these representations are organized remain controversial. Specifically-it is still debated whether the neural responses of high-level visual cortex are organized according to visual, pictorial properties or along semantic, abstract information. This question reflects on the general issue of cortical domain specificity and modularity and consequently has been the subject of extensive discussion (for a recent review, see^10^).

Recent investigations using deep learning algorithms offer new tools to explore this question (for recent perspectives supporting this approach see ^11–13^). In the past decade, deep learning algorithms have reached human level performance in object and face recognition. Moreover, the relational structure of their embeddings (which will be termed here-*representation*) generated by these algorithms across their different layers are correlated with the representation of the neural response to the same images in early and high-level areas of visual cortex^12,14–17^. Intriguingly, recent studies show that the neural response to visual stimuli can be predicted not only by visual deep learning algorithms but also by the linguistic representation of their textual descriptions by large language models (LLMs)^18–21^. This surprising finding has been suggested to support the notion that the representations of images in high-level visual brain areas are aligned with semantic information^19,22,23^.

In particular, Conwell et al (2023) used visual and language deep learning algorithms to predict the fMRI response of participants viewing images from the natural scene dataset^24^. In this dataset, each image is accompanied by verbal descriptions of the content of the image (e.g., “A horse carrying a large load of hay and two people sitting on it”). This enabled them to ask whether the neural response to the images can be predicted not only by the visual representation of the images based on visual deep convolutional neural networks (DCNNs), but also by LLMs’ embedding of the verbal descriptions of the images. Results showed that the response of the occipito-temporal cortex to images was similarly predicted by the visual representations of visual DNNs as well as the linguistic representations of the LLMs^20^. Using the same dataset, Doerig and colleagues (2024) used a searchlight analysis on the fMRI - derived maps and found that LLM representations accounted for brain responses to images across the entire visual system, particularly in higher-level areas^19^. Additionally, they showed that using text descriptions about people, scenes, or places, as input to their LLM brain encoder, predicted the response of brain areas that are visually selective to these categories (e.g., sentences about people/places/food were correlated with face/place/food-selective ROIs, respectively). Accordingly, they concluded that the ventro-temporal visual cortex represents semantic information about visual input. They further suggested an alignment between visual and linguistic representations in predicting the response of high-level visual cortex to images.

However, the textual descriptions used in these studies^18–20^ were largely limited to the visual content of the images. Yet, visual images can also be described using textual information that is independent of their visual content, conveying conceptual or abstract knowledge acquired about them. Thus, it remains unclear whether the visual-language alignment observed in high-level visual cortex may be restricted to visual text or could extend also to abstract textual descriptions that are independent of the visual content of the images. In the present study we distinguish between two types of linguistic descriptions of visual images: a linguistic description of the image’s purely visual content, which we term *visual text*, and a linguistic description of conceptual knowledge that cannot be derived from the immediate visual attributes of the image (such as the name of the profession or the geographical location of a familiar person/place), which we term *abstract text* (see Fig 2A for an example). The first represents pictorial visual information coded in a linguistic form, whereas the second represents conceptual/abstract knowledge that cannot be derived from the image’s immediate visual content and necessitates further linguistic knowledge^25^. We hypothesize that *visual text*, but not *abstract text*, will predict the correlation of neural responses in high-level visual cortex and LLM embeddings.

Our hypothesis is partially supported by a recent iEEG study ^26^ that used visual DCNN representations and linguistic LLM representations to predict the neural responses to familiar faces and places. In this study, the linguistic descriptions were based on Wikipedia entries for the familiar stimuli (see Fig. 2A), which primarily contained conceptual information unrelated to the visual content of the images. The results showed a strong correlation between neural responses to images and visual DCNN representations but a weak correlation with LLM representations of abstract text^26^. Here we hypothesize that the linguistic form of visual information is aligned with visual content of the images, whereas the abstract textual description is likely represented in fronto-parietal visual areas. To test this hypothesis, both visual and abstract textual descriptions should be generated for the same images and used to predict their neural responses in the visual cortex.

To that end, we used visual and language deep learning algorithms to predict the iEEG responses obtained from high order visual areas in a unique data set containing images of familiar faces or places (Figure 1). Familiar faces and places are advantageous to test our hypothesis as they enable us to generate abstract descriptions that are independent of the visual appearance of the image. We generated two types of textual descriptions, *visual text*, which describes content that can be derived from visual information in the image, and *abstract text*, based on the Wikipedia definition of the familiar image, which is independent of its visual content (see Figure 2; see supplementary material for visual and abstract textual descriptions of each image). We hypothesized that the response of high-level visual cortex to the familiar images would be aligned with LLM representations of their visual textual descriptions, replicating previous findings^19,24^, but not with the LLM representations of abstract textual descriptions of the same images, in line with^26^. To further validate the visual and conceptual nature of the DNNs representations, we also collected human similarity ratings for the same set of images. We asked human participants to rate the visual similarity of the same images, or the semantic similarity of their name labels and examined the correlations of these ratings with the visual and linguistic DNN representations. To extend these findings beyond the limited set of stimuli used in the iEEG study, we replicated the similarity rating study with another set of visual images.

**Figure 1:**
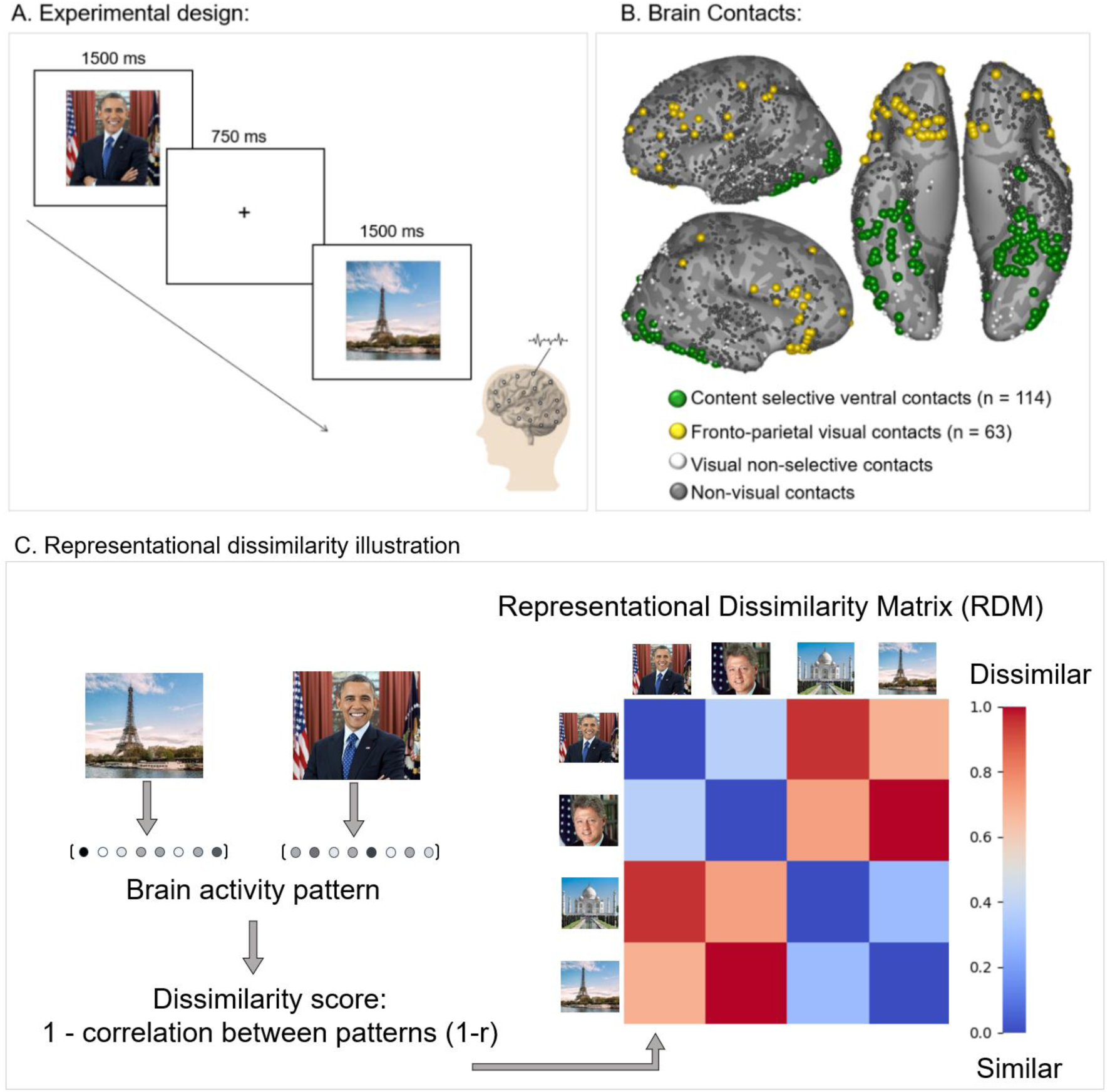
Experimental design: **A.** Participants viewed 28 images of familiar faces and places while their neuronal activity was recorded via intracranial EEG. **B.** Contact locations: yellow - Fronto-parietal visual contacts, green – Content-selective ventro-temporal contacts, white – visual non-selective contacts, gray-non-visual contacts. **C.** illustration of brain representational dissimilarity scores calculation

**Figure 2:**
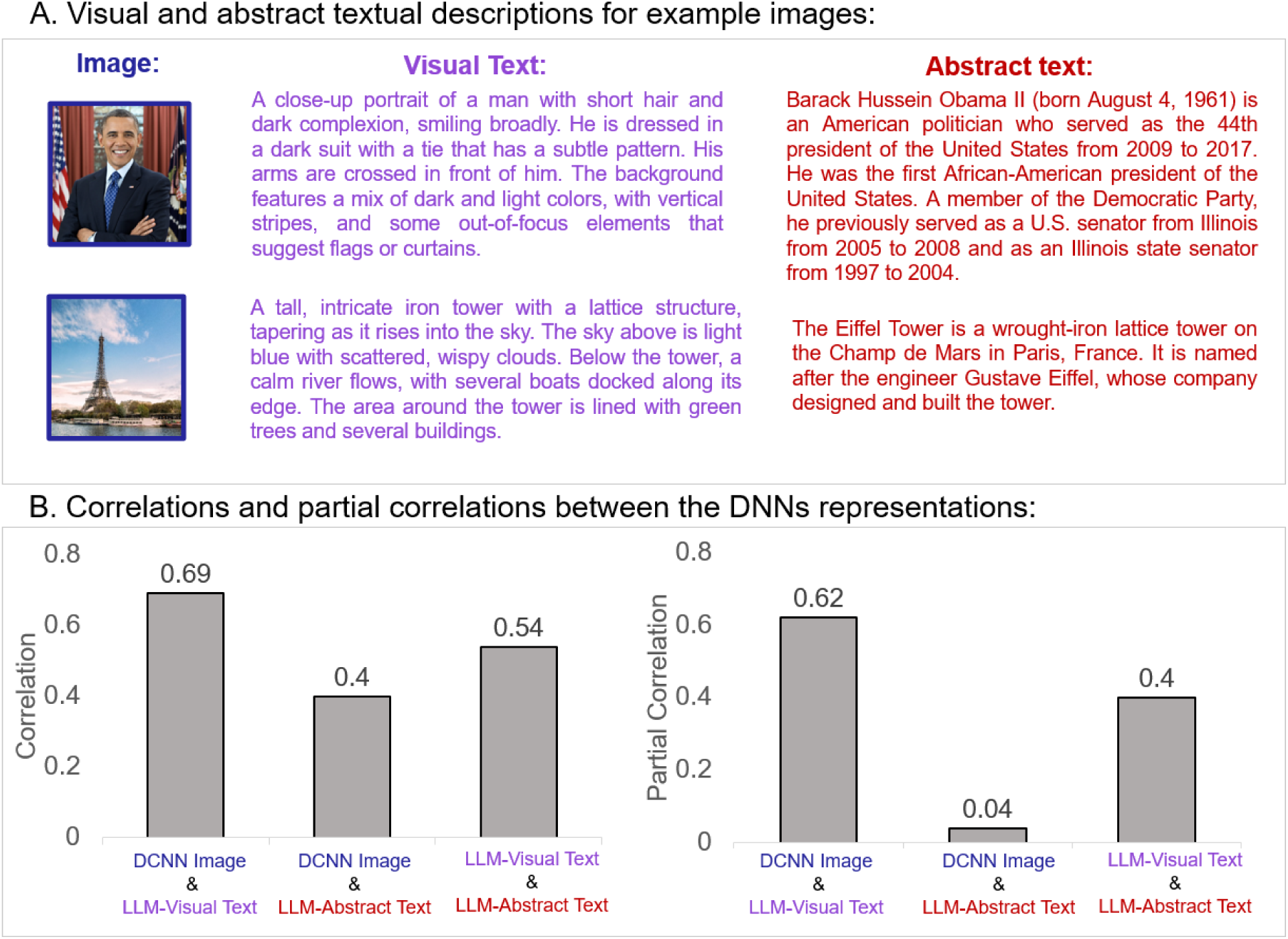
Examples of two images and their visual- and abstract-textual descriptions, and the partial correlations between their visual and language DNNs representations: **A**. Examples of images and their visual (purple) and abstract (red) textual descriptions: **B.** A pre-trained object DCNN (VGG-16) was used to extract the embeddings of the images (DCNN-image), and a large language model (GPT-2) was used to extract the embeddings of the visual and abstract textual descriptions. The correlations and partial correlations between the RDMs of the DNNs were high between images and visual text but low between images and the abstract text. All images in the figure were replaced by licensed images with a similar appearance from Freepik (https://freepick.com).

## Results

### Experimental design and intra-cranial contacts information

Our study involved intra-cranial recordings, conducted for clinical purposes in 13 patients (10 females, mean age 34.7 ± 9.6) covering 2571 recording sites. During the experiment the participants viewed 28 images of well-known individuals and places (14 of each category), Each image was presented 4 times in semi-random order (for further details see Methods and ^27,28^). It should be noted that the IEEG method, while providing superior spatio-temporal resolution of the neuronal signals, imposes strict limits on the duration of the experiments-hence the limited number and categories of visual images that could be used in the study.

Figure 1 shows the experimental design (Fig. 1A) and recording sites (Fig. 1B) that were examined in the present study. First, we identified the visually responsive contacts (n=377, out of the 2571 total contacts) and then categorized them based on their anatomical locations (ventro-temporal, fronto-parietal) and functional characteristics, into different regions of interest (ROIs) (see methods of ^27^ for details). To test our hypothesis, we focused on the ventro-temporal visual ROI and used the fronto-parietal visual contacts as a control ROI.

### The relational structure of the DNN representation of visual images is correlated with visual text but not abstract text

For each of the familiar images we generated two types of textual descriptions: one that describes pictorial information that can be derived directly from the visual content of the image (visual text) (see Methods for further details) and another, abstract description, was obtained through their Wikipedia definition (abstract text). Careful reading of the Wikipedia-based texts verified their strong bias towards abstract meaning that cannot be derived from their visual appearance alone (See Figure 2A, and full set of textual descriptions in the Supplementary material). To assure that the textual descriptions are uniquely associated with each of the images we asked participants to match each text to one of the images from the same category. Both text types were strongly associated with the images: for visual descriptions, the text was correctly matched by an average of 93.6% of participants (mode = 100%, minimum = 70%), and for abstract descriptions, by an average of 95% (mode = 100%, minimum = 80%). We then used visual and language DNNs to construct the visual and abstract representation of the stimuli, respectively. The representations of the images were derived from the final fully connected layer of an ImageNet trained DNN (VGG-16^29^). The representations of the visual and abstract text were extracted using the final layer of GPT2^30^ (See Methods section for details).

We then generated a representational dissimilarity matrix (RDM) for each representation, by computing the cosine distance between the embeddings of the 28 images/textual descriptions for each input. This resulted in three different DNN-generated RDMs, one image-based (DCNN-Images), one visual-text based (LLM-Visual), and one abstract-text based (LLM-abstract). We first asked, by comparing the RDMs of the different DNNs, if the DNN-generated representations of the images are more similar to their DNN-generated visual-text representations compared to the representation generated by the abstract text descriptions. The zero order correlation between the representation of the images with the visual text (r=0.69) was significantly higher than the correlation between the representations of the images with the abstract text (r=0.41), and the correlations between the two types of texts (r=0.55). To assess the unique association between each pair of DNNs, independent of their shared variance with the third DNN, we calculated the partial correlations between each pair of DNNs representations when the third DNN is held constant. As shown in Figure 2B, the partial correlation between the images and the abstract text (when removing their correlation with the visual text) is not significant (0.04), suggesting that the abstract textual descriptions are independent of the visual content of the images. The partial correlation of the visual text was higher with the images (0.61) than with the abstract text (0.41). These findings demonstrate a significant correspondence, in the artificial networks, between the representations of images and their visual textual descriptions, but no association between the representations of images and their abstract textual descriptions.

### Visual ventro-temporal cortex is highly correlated with DNN’s image-based and visual-text based representations, but not with abstract-text based representation

To examine the similarity between the organization of the three representations generated by DNNs and the representation of high order ventro-temporal visual cortex of the same images, we created a representational dissimilarity matrix (RDM) based on the iEEG responses for the different stimuli. For each stimulus, we calculated the average neural response (amplitude of the broad band high frequencies of the iEEG signal in each contact ^28^) over the four stimulus repetitions, within the 0.1 to 0.4 second window after stimulus onset, in each contact in the ROI. We selected this time window because it captures the peak neural visual response^28^. We used the averaged iEEG derived population vector as the response pattern of each stimulus within the ROI. We then measured the Pearson correlations between the activation patterns of each pair of stimuli. The dissimilarity between the neuronal representations of two stimuli was represented as one minus the Pearson correlation value (thus lower values indicate greater similarity).

Next, we compared the relational-structure (i.e. RDM matrices) similarities between the response of the visual ventro-temporal cortex to the images, and the three artificial representations generated by the DNNs for the same stimuli seen by the patients. The similarities between the brain-based and DNN-based relational structures were computed as the Pearson correlation between the brain and the respective DNN-derived RDMs. To test for statistical significance, we computed the same correlations using a permutation test (shuffled pairs labels, see methods for further details). We tested both the significance of each predictors’ (DNNs) correlation individually and the difference between each correlation in each ROI. P-values were FDR corrected for each ROI and each analysis (individual predictors and differences) separately.

Our analysis shows that the neural response of the visual ventro-temporal cortex was similarly correlated with the visual representation of the image (DCNN-Image), and the visual-text representation (LLM-visual) but showed a significantly lower correlation with the abstract text representation (LLM-abstract) (Fig. 3 left top panel). The same analysis with the responses of the fronto-parietal contacts to the images showed significant correlations with all three DNN-based representations, but no significant difference between them. To examine the unique similarity of each DNN representation to the response of the ventro-temporal visual contacts to the images, we calculated the partial correlations between each DNN-based RDM and the RDM derived from the visual cortex, when the two other DNN representations are held constant. The same correlational analysis was performed on the response of the fronto-parietal contacts.

**Figure 3:**
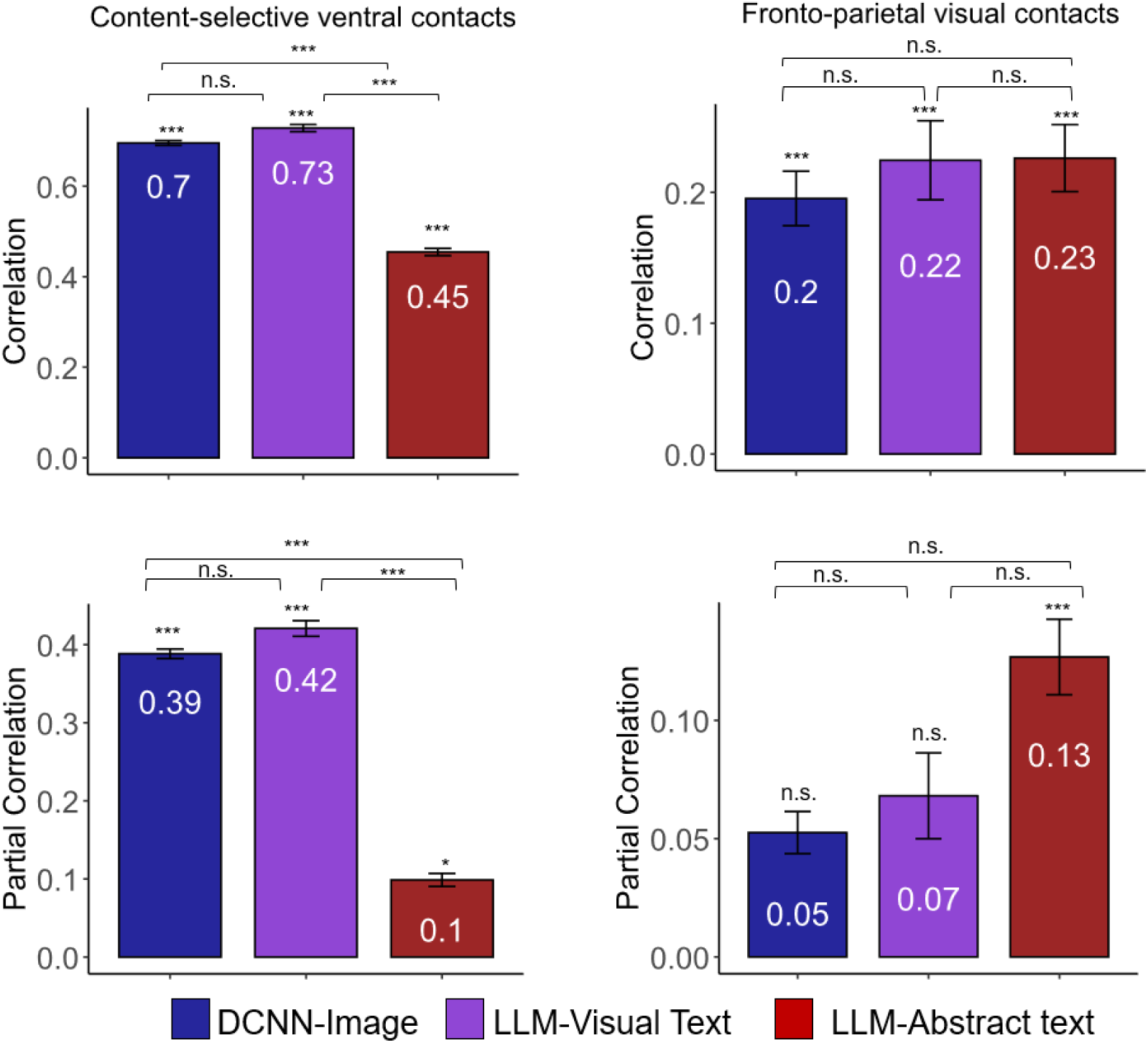
Correlations between the representations of ventro-temporal content-selective and fronto-parietal visual contacts to images and the representations of the images by: visual DNN (DCNN-image), visual textual descriptions (LMM-visual text), and abstract textual descriptions (LMM-abstract text): Zero order correlations of each DNN RDM’s with RDMs from brain ROIs are presented in top panels; partial correlations with the RDMs of brain activity patterns, when the other two DNNs are held constant, are presented in bottom panels. Left: the correlations and partial correlations with the Content selective ventral contacts (n=114). Right: correlations with the Fronto-parietal visual contacts (n=63), Error bars indicate leave one participant out procedure s.e.m. All *p* values were derived from a pair-images two-sided permutation test (10000 permutations). Reported *p* values are FDR corrected in each ROI separately. Note the clear drop in correlation from visual DCNN and LLM-Visual to LLM-Semantic in the ventral contacts. ∗ *p*_*FDR*_ < 0.05, ∗∗ *p*_*FDR*_ < 0.01, ∗∗∗ *p*_*FDR*_ < 0.001

Both the image-based (DCNN-Image) and the visual-text based (LLM-Visual) representations uniquely predicted the ventro-temporal visual cortex response to the images, whereas the abstract text did not account for any additional information. There was no significant difference between the correlations of the image and the visual-text, and they were both significantly higher than the abstract text (Fig. 3 left bottom panel). The response of the fronto-parietal visual contacts showed the reverse pattern, where only the abstract-text representation significantly accounted for unique information in their neural response (Fig. 3 right bottom panel).

### Temporal dynamics show that over time the response of the visual ventro-temporal cortex is mainly explained by the image representation

The iEEG recordings enable us to also examine the temporal dynamics of the visual and linguistic representations. We ran the same correlational and partial correlation analysis of the DNNs with the neural responses across time (see Figure 4 panels A and B). Findings show that correlations with the visual DNN representation persists for longer duration until stimulus offset, whereas the DNN representation of the visual text is short lasting (see Figure 4 panel B).

**Figure 4:**
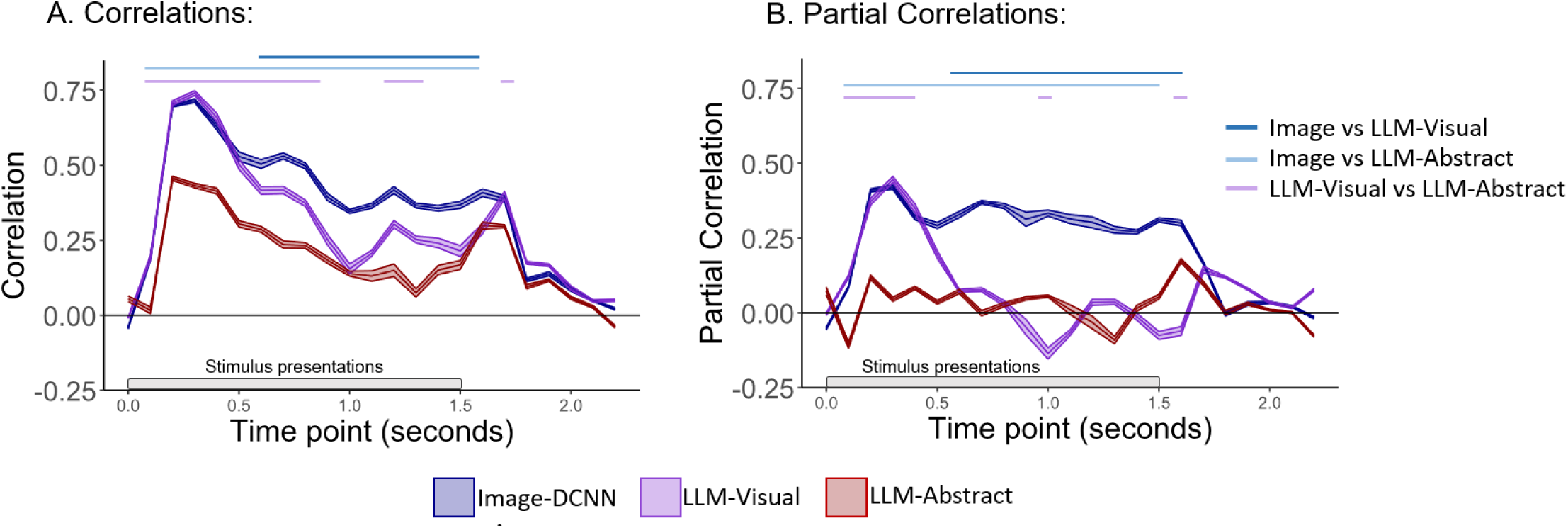
Correlations and partial correlations over time, between the representations of ventral content-selective contacts to images and the representations of the images by: visual DNN (DCNN-image), visual textual descriptions (LMM-visual text), and abstract extual descriptions (LMM-abstract text): **A.** Zero order correlations of each DNN RDM’s with he RDM of the ventral content-selective contacts’ activity (n=114) **B.** Partial correlations with the RDM of brain activity patterns, when the other two DNNs are held constant. Ribbon indicates leave one participant out procedure s.e.m. The lines above represent significance, all *p* values were derived from a pair-images two-sided permutation test (10000 permutations).

### Visual-text representation is more correlated with visual than semantic similarity ratings

To further examine if the visual and abstract textual descriptions represent visual or semantic information about the images, we asked human participants to rate the visual or semantic similarity of all possible pairs of the 28 stimuli. One group of participants (n = 20) was asked to rate the visual similarity between each pair of images based on their visual appearance. Another group of participants (n = 20) was presented with the names of the stimuli (the name of the identity or place presented in the image) and was asked to rate the semantic similarity between each pair of items based on the conceptual knowledge they had about them (see Fig. 5A; see Methods for further details). For each participant we created an RDM of the stimuli, based on the participant’s similarity ratings. To examine the unique contribution of each of these representations, we computed their partial correlations. We performed this analysis once when including only the DCNN-Image and the abstract-text representations, and once when including only the visual-text and the abstract-text representations.

**Figure 5:**
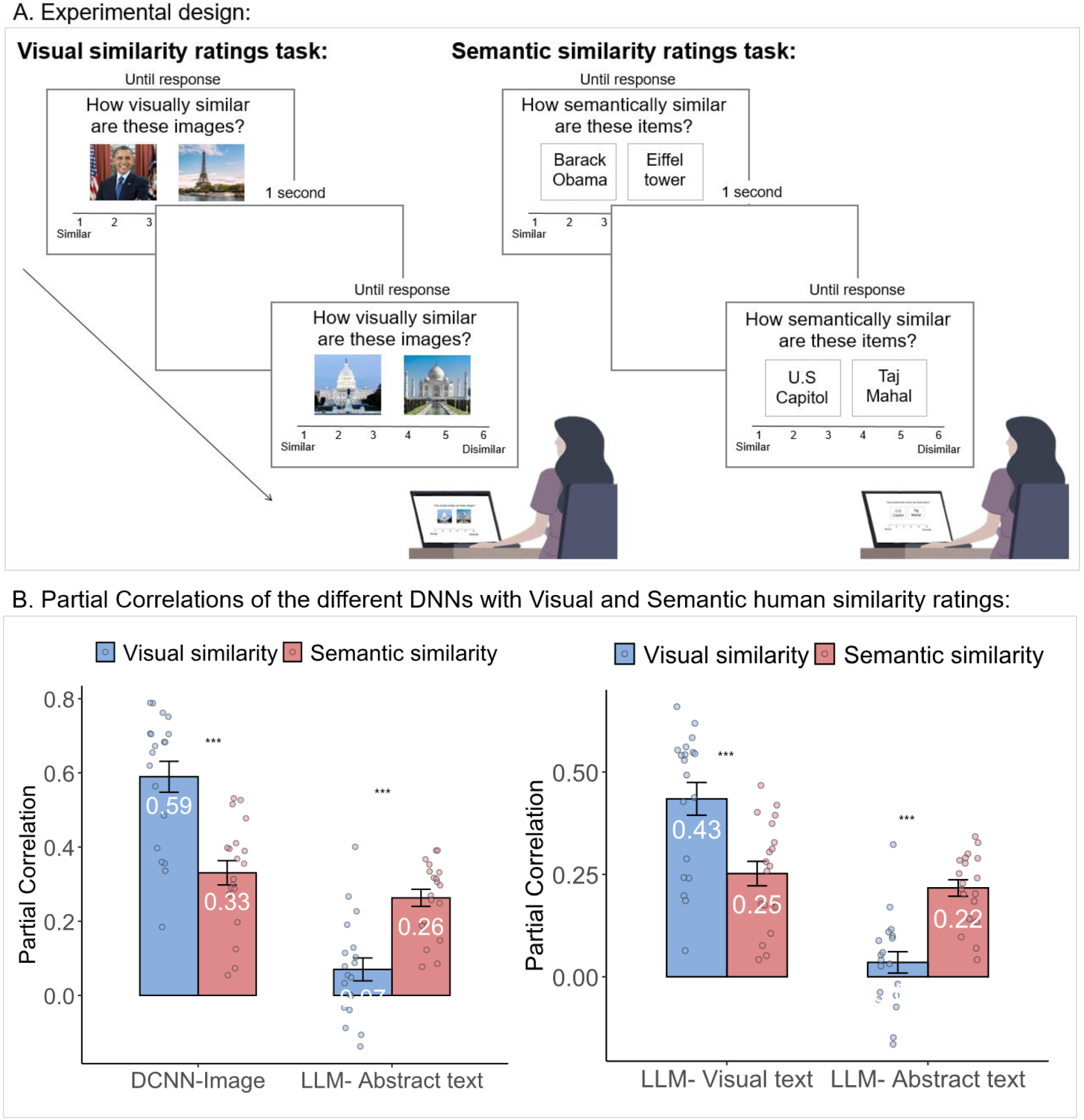
Experiment design and results of human similarity ratings: ***A.*** Experimental design: Participants rated the visual similarity (n = 20) of famous faces and places while presented with their images that were used in the iEEG study (left) or the semantic similarity (n = 20) between the names of the same stimuli presented in the images (right) ***B.*** Partial correlations with human similarity ratings: Left panel-When including only the DCNN-Image and the abstract-text representations right panel-When including only the visual-text and the abstract-text representations. Both figures depict significant interaction and similar results. Error bars are s.e.m. between participants. p values were derived from a post hoc compression between task and predictors. Reported p values are FDR corrected.∗∗ *p*_*FDR*_ < 0.01 ∗∗∗ *p*_*FDR*_ < 0.001. All images in the figure were replaced by licensed images with a similar appearance from Freepik (https://freepick.com).

As shown in Fig. 5, these two analyses resulted in a very similar pattern of results with the representation of the visual and visual-text DNNs resulting in higher partial correlation with human visual than semantic similarity rating, whereas the representation of the abstract-text showed the reversed pattern. To test the statistical significance of these findings, the partial correlations were transformed to fisher z-scores (the figures show the original partial correlation before fisher-z transformation) and a mixed ANOVA was performed with DNN-representation (image and abstract-text or visual-text and abstract-text, respectively) as within-participant factor, and task (visual similarity rating, semantic similarity rating) as between-participants factor, on the Fisher transformed partial correlations. The ANOVA including DCNN-Image and abstract-text representations revealed a significant main effect for DNN (F(1, 36) = 75.7, p < 0.001, η_p_^2^ = 0.68) and for Task (F(1,36) = 4.13, p = 0.049, η_p_^2^ = 0.10), and a significant interaction between them F(1, 36) = 46.24, p < 0.001, η_p_^2^ = 0.56). Post-hoc comparison between visual and semantic rating tasks for DNN-based representation (FDR corrected) revealed that image representations show higher partial correlation with visual than semantic similarity ratings (t (72) = 6.29, p < 0.001), but abstract-textual representations show higher partial correlation with the semantic rather than the visual rating tasks (t (72) = 3.44, p < 0.001). The ANOVA including visual-text and abstract-text representations revealed a significant main effect for DNN (F(1, 36) = 43.8, p < 0.001, η_p_^2^ = 0.55) but not for Task (F(1,36) = 0.33, p = 0.57, η_p_^2^ = 0.009), and a significant interaction between them F(1, 36) = 30.54, p < 0.001, η_p_^2^ = 0.46). Post-hoc comparison between visual and semantic rating tasks for DNN-based representation (FDR corrected) revealed that visual-text representations show higher partial correlation with visual than semantic similarity ratings (t (72) = 4.64, p < 0.001), but abstract-textual representations show higher partial correlation with the semantic rather than the visual rating tasks (t (72) = 3.92, p < 0.001).

These results are in line with our conclusion that, similar to the visual-image representation, visual-text representations are aligned with visual rather than semantic information.

### Using a different set of stimuli revealed a similar pattern of correlations between the DNNs and the human similarity ratings

As mentioned above, visual and abstract texts can be clearly dissociated for familiar images, as our acquaintance with them involves acquiring non-visual verbal knowledge. However, to extend our findings beyond familiar images, we also ran the same visual and semantic similarity task with unfamiliar scenes. This also allowed us to overcome a shortcoming of the current study in which, due to the constraint imposed by the unique clinical conditions of the IEEG recordings, the number and types of images that we used were limited. It was therefore important to validate the relationship between visual text and abstract text to visual and semantic representations with an additional set of images. To that end, we used 15 images of different indoor and outdoor scenes taken from Bainbridge et al^31^. We extracted visual and abstract descriptions of the images (see Figure 6A for an example of a kitchen and a playground) as we did for the first set of images. Then we used a DCNN to extract the images representations and an LLM to extract the representation of both the visual and the abstract textual descriptions (see methods for further details). Figure 6B shows that the correlation and partial correlation of the images are much higher with visual text than with abstract text representations. We asked participants to make visual or semantic similarity ratings of all possible pairs of the 15 stimuli belonging to the new set. As with the original set of images, one group of participants (n = 20) was asked to rate the visual similarity between each pair of images based on their visual appearance. Another group of participants (n = 20) was presented with the names of the stimuli (the name of the scene presented in the image, for example “Kitchen”) and was asked to rate the semantic similarity between each pair of items based on the conceptual knowledge they had about them (see Methods for further details). Then we calculated the partial correlation between the participants’ visual and semantic similarity ratings with the DNNs representations, between DNN pairs, holding one DNN constant in each calculation. The results, as shown in Figure 6C, were very similar to the first set of stimuli (Figure 5B). The partial correlation used for the analysis were transformed to fisher z-scores (the figures show the original partial correlation before fisher-z transformation).

**Figure 6:**
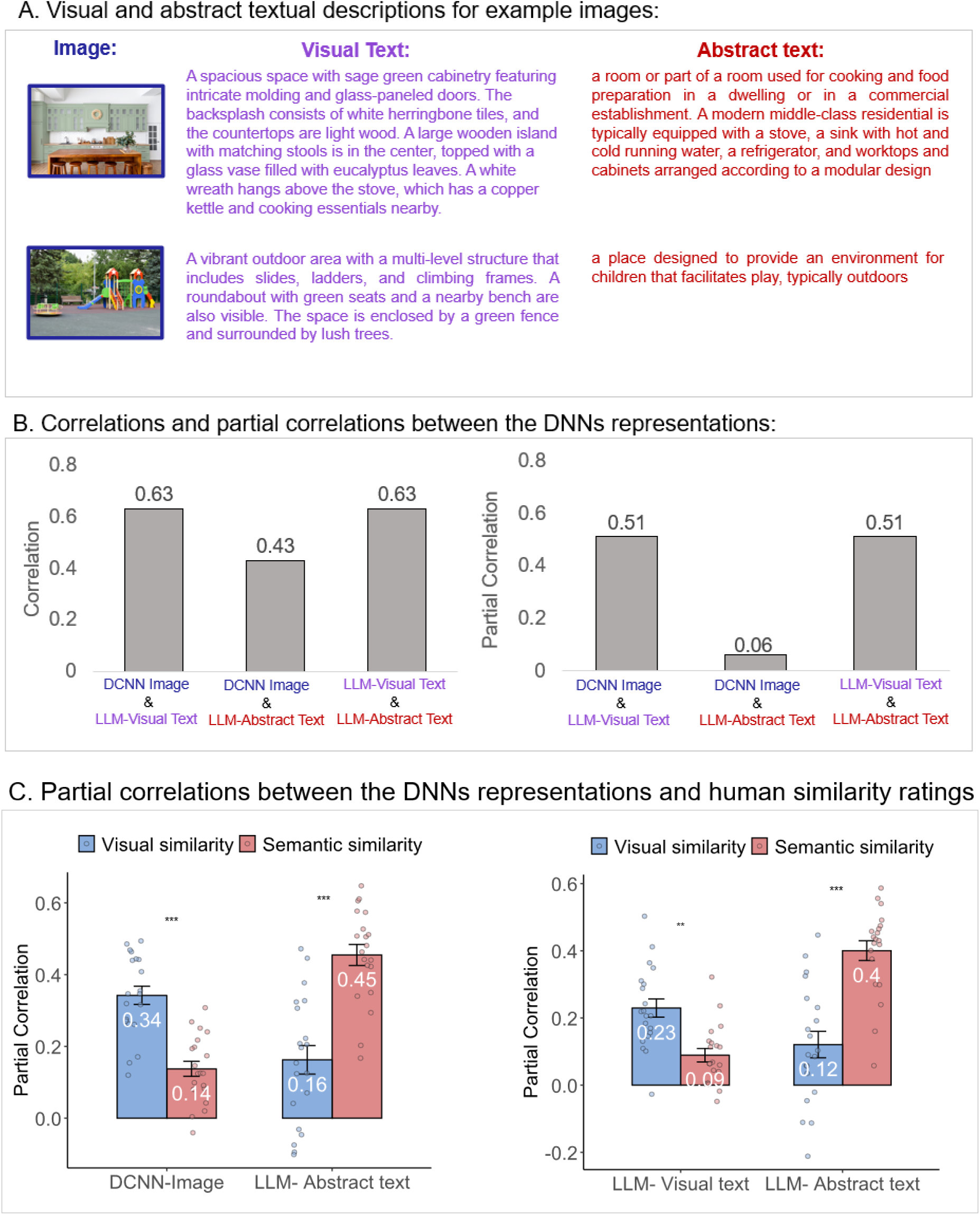
The results using a different set of images: **A.** Examples of two scene images, a kitchen and a playground, and their visual (purple) and abstract (red) textual descriptions: **B.** A pre-trained object DCNN (VGG-16) was used to extract the embeddings of the images (DCNN-image), and a large language model (GPT-2) was used to extract the embeddings of the visual and abstract textual descriptions. The correlations and partial correlations between the RDMs of the DNNs were high between images and visual text but low between images and the abstract text. **C.** Partial correlations with human visual and semantic similarity ratings when including only the DCNN-Image and the abstract-text representations (left) or when including only the visual-text and the abstract-text representations (right). Both figures in panel c depict significant interaction and similar results. Error bars are s.e.m. between participants. *p* values were derived from a post hoc compression between task and predictors. Reported *p* values are FDR corrected.∗∗ *p*_*FDR*_ < 0.01 ∗∗∗ *p*_*FDR*_ < 0.001. All images in the figure were replaced by licensed images with a similar appearance from Freepik (https://freepick.com).

The ANOVA including DCNN-Image and abstract-text representations revealed a significant main effect for DNN (F(1, 38) = 4.85, p = 0.034, η_p_^2^ = 0.11) and for Task (F(1,38) = 4.29, p = 0.045, η_p_^2^ = 0.10), and a significant interaction between them F(1, 38) = 52.54, p < 0.001, η_p_^2^ = 0.58). Post-hoc comparison between the visual and semantic rating tasks for DNN-based representation (FDR corrected) revealed that image representations show higher partial correlation with visual than semantic similarity ratings (t (76) = 4.74, p < 0.001), but abstract-textual representations higher partial correlation with the semantic rather than the visual rating tasks (t (76) = 7.09, p < 0.001). The ANOVA including visual-text and abstract-text representations revealed a significant main effect for DNN (F(1, 38) = 8.5, p = 0.006, η_p_^2^ = between them F(1, 38) = 33.34, p < 0.001, η_p_^2^ = 0.46). Post-hoc comparison between visual and semantic rating tasks for DNN-based representation (FDR corrected) revealed that visual-text representations show a higher partial correlation with visual than semantic similarity ratings (t (76) = 3.24, p = 0.001), but abstract-textual representations display a higher partial correlation with the semantic rather than the visual rating tasks (t (76) = 6.75, p < 0.001). These results further support our conclusion that visual-text representations are aligned with visual rather than semantic information, whereas abstract text representations are correlated with semantic rather than visual information.

## Discussion

Many neuroimaging and IEEG studies have convincingly shown that the representations generated for images by DCNNs trained for recognizing images are strongly correlated with neural responses in the high-level visual cortex^4,14–16,32^. Interestingly, recent fMRI studies reveal that linguistic representations of textual descriptions generated by large language models also achieve similarly high predictions of the neural response to images in high level visual cortex^19–21^. These findings suggest a strong alignment between visual and linguistic representations. However, the question of the nature of this linguistic information was largely left unaddressed. Does it reflect purely visual and pictorial aspects of the images or also more abstract, conceptual knowledge, that cannot be derived directly from the visual images themselves? This is an important question, as it determines the interpretation of this intriguing finding, regarding the information guiding the organization of high-level visual cortex. To address this question, we generated two types of linguistic representations to the same images, one that describes information that can be extracted directly from the images and the other that is independent of the pictorial visual content of the image and conveys abstract knowledge about it that can only be gained through non-visual channels (Figure 2). We hypothesized that only the former is aligned with neural responses in visual cortex. To test this hypothesis, we employed direct iEEG recordings of neural responses to images of famous faces and places, obtained during clinical diagnostic procedure in epilepsy patients (see methods). In this type of stimuli, abstract information, such as the name or occupation of the person or the name and location of a place, is independent of the visual appearance of a specific image (see Fig 2A and Supplementary Table 1). We then used a visual DCNN to generate an image-based representation, and a LLM to represent the visual and abstract textual descriptions; next, we tested the alignment of each with the high-level visual cortex representations. Our findings show that DNN representations of images and of visual-text similarly predicted the neural response of high-level visual cortex to images, consistent with recent reports^19–21^, whereas abstract textual representations showed significantly lower correlations and did not explain additional variance beyond the visual information. These findings support our hypothesis that the alignment between linguistic information and the high-level visual cortex response to images, is restricted to textual descriptions of pictorial, visual information. It further validates the important distinction between visual and abstract linguistic representations.

To further examine if the visual, visual-textual and abstract-textual DNN representations encode the subjects’ visual or semantic perceived information, we asked human participants to rate the visual or semantic similarity of the same images. Our findings show that visual-text was more correlated with visual similarity ratings than with semantic similarity ratings (Figures 5 and 6). This pattern of correlation corresponds to the pattern of correlation in high-level visual cortex (Figure 1B), and further emphasizes the visual nature of this representation. This same pattern of findings was found for another set of images (Figure 6), supporting the notion that results that were found for the images used in the iEEG study are not specific to this image-set.

Our findings that visual textual representations are aligned with the neural response to images are consistent with recent findings that show that linguistic descriptions of scenes predict the neural response of high-level visual cortex to the corresponding images^18–21^. These results were even replicated in a monkey fMRI study that used the same stimuli, further indicating that these representations do not reflect the operation of a language system^18^. These verbal descriptions depict categorical information that can be extracted from visual inspection of the image.

The question of whether high-level visual cortex represents visual features or semantic categories has been extensively debated (for reviews see ^3,10^). Several studies have suggested that low level visual features such as eccentricity, spatial frequency^33^ aspect ratio^34^ or curvature^35^ underlie the large-scale functional organization of high-level visual cortex. A main challenge in dissociating between visual features and semantic categories is that they are not orthogonal. For example, most fruits are round or curvy whereas most tools are long and straight. To disentangle between shape and category, Bracci and Op De Beeck (2016)^36^ measured the fMRI response to objects from different categories (e.g., animate, musical instruments, tools, food) that are matched as much as possible in shape. They found that the organization of the visual cortex transitions from shape, in posterior low level visual areas, to category representations, in lateral and ventro-temporal visual brain areas. They further conjecture that selectivity to visual features in these regions may primarily serve a possible goal of the system to classify images to different categories.

While our iEEG study emphasizes visual over semantic organization principles in the local organization of high order visual areas, this finding does not necessarily contradict a role for such information at coarser, large scale, levels e.g. those defining major cortical subdivisions such as revealed in fMRI studies. For example, in an fMRI study, Carlson et al (2014)^22^ showed that word co-occurrences and behavioral semantic similarity judgments predicted the representational geometry of the neural response to the coarse distinction between cortical areas preferring animate vs. inanimate categories in the inferior temporal cortex. Popham and colleagues (2021)^23^ extended these findings by showing that for each category-selective visual ROI in the visual cortex, there is an adjacent category-selective semantic ROI for the same category, suggesting visual-semantic alignment at the level of category selective cortical areas. However, it is important to emphasize that such large scale studies do not address the local relational structures within each individual areas-which show exemplar specificity within a category (e.g faces ^16,26^). For such detailed analysis, higher spatial resolution methods such as iEEG are especially informative^37^.

The partial correlation analysis further shows that visual and visual-textual information contributed distinct proportion of variance in the neural response to images. Analysis of the temporal dynamics of the representation also indicates that the visual representation shows higher correlation for longer duration till stimulus offset, whereas the visual linguistic representation is short lived. This suggests that in addition to their aligned representations the neural responses also capture different aspects of the information contained in the images. Thus, our findings suggest that there is additional unique information that visual and linguistic representations convey about the visual content of an image beyond their aligned representations. This finding is relevant to the question of whether sensory and linguistic information about the physical world reflect overlapped or distinct types of information, which has been highly debated. Embodied cognition theories posit that all information is grounded in sensory information^38,39^, whereas dual-code models suggest a distinction between visual and language inputs^40^. The findings that the representational geometry of color in congenitally blind individuals is very similar to the representation of sighted individuals in specific parietal cortical regions demonstrates that visually-aligned linguistic representations can be generated in the absence of visual input in specialized cortical regions^40^.

The distinction between visual and abstract linguistic information may correspond to the distinction between concrete/grounded vs. abstract words, respectively, which has been extensively studied for several decades in the language domain^41,42^. These studies reveal substantial cognitive and neural differences between the representations of concrete/grounded words, such as names of objects as opposed to abstract words, such as justice and love. However, the comparison between concrete and abstract words inherently involves comparing very different concepts, whereas the visual and abstract textual descriptions used in the current study are linguistic descriptions of the same stimulus. Future studies that concern the difference between grounded and abstract linguistic information ^43–45^ may use our approach to compare the neural representations of these two types of verbal information that describe the same concept.

In conclusion, our study supports the notion that the alignment between the neural response of high-level visual cortex to images and linguistic representations is limited to pictorial visual information presented in a linguistic code. Specifically, we demonstrate a clear neural distinction between visual-textual and abstract-textual representations. Visual-textual information was strongly aligned with neural relational structures in high-level visual cortex, whereas abstract-textual information was not. These findings demonstrate the specialized role of the visual cortex in processing information directly derived from visual stimuli.

## Materials and Methods

### iEEG study Familiar faces and places

#### Stimuli

28 color images of famous identities and famous places, 14 of each category. See Supplementary Material for the list of all identities and places. During the iEEG recordings the participants did not indicate whether they were familiar with the identities and places before the experiment. Therefore, to ensure that the images used are indeed generally familiar items, in a different study we asked 40 participants to indicate whether they are familiar with each face or place. Across all participants, the images were recognized by an average of 90% (mode = 97%), confirming that these faces and places were highly familiar stimuli.

#### iEEG data: experimental design and intra-cranial contacts information

We describe here the participants’ tasks and stimuli as well as ROI definitions. Additional details can be found in ^27^.

##### Participants

Electrophysiological data from 13 patients (10 females, mean age 34.7 ± 9.6) were collected via intracranial EEG during pre-surgical evaluation for drug-resistant epilepsy at North Shore University Hospital, NY. As part of the clinical assessment, subdural or depth contacts were implanted in all patients. All patients gave fully informed consent, including consent for publication of the results. No clinical seizures occurred during the experiment. The Feinstein Institute for Medical Research’s IRB review board monitored the study, and the study followed the latest declaration of Helsinki and US National Institute of Health guidelines.

##### Experimental task and stimuli

The experiment was composed of two runs in which the participants viewed 28 images of famous faces and places, 14 images of each category. Each run started with a 200-second resting phase with closed eyes. Then, the participants were presented with the images, each image was presented for 1500 ms, with 750 ms inter-stimulus intervals. In each run, the participants saw 14 different images (7 of each category), and each image was presented four times in a semi-random order to avoid consecutive repeats. The participants were instructed to memorize the details of the images, as they will be required to recall the images and describe their visual features in a later recall phase. The recall phase is not analyzed in this study. It is noted that the participants performed well, and no participants were excluded based on performance (Further details can be found in ^27^).

##### Definition and grouping of visually responsive contacts

The analyses in this study were conducted only on visually responsive recording sites (contacts), subdivided into different ROIs. Therefore, we extracted out of all 2571 recording sites the visually responsive contacts, which we defined as electrodes displaying statistically significant activations in response to the images. The data from all 2571 recording sites was preprocessed and transformed to the high-frequency broadband (HFB) signal (see ^28^ for further details). We averaged the HFB response across the 100 to 500 ms period following stimulus onset and compared each contact’s post-stimulus response to the baseline response prior to stimulus onset. The pre-stimulus baseline was then averaged across the - 400ms to −100ms interval prior to stimulus onset. We compared the stimuli-evoked response to the baseline response, using a two-tailed Wilcoxon signed-rank test, that was FDR corrected for all contacts (from all patients) together. We defined visually responsive contacts as contacts demonstrating a significant HFB difference (*p*_*fdr*_ < 0.05), resulting in 377 visual contacts.

Then we divided the visually responsive contacts based on anatomical and functional parameters, to content-selective ventro-temporal and fronto-parietal visual contacts. To define the content selective contacts, we examined the visually responsive contacts in six anatomical regions across the ventro-temporal visual hierarchy, based on the Desikan Killiany atlas. These regions included the lateral occipital cortex (LO), inferior temporal gyrus (ITG), lingual gyrus, parahippocampal gyrus (PHG), fusiform gyrus, and entorhinal gyrus, excluding early visual contacts. Then, for each contact, we calculated the averaged HFB response to each stimulus during the 100-500ms post-stimulus window. Contacts in the selected regions that showed significant content selectivity, defined as a difference of at least 3.5 SD between the top 10 preferred images (averaged response) and bottom 10 images (averaged response), were included in the content-selective contact set (n = 114). We defined the fronto-parietal ROI based on the Desikan Killiany atlas locations of the following labels: superior frontal gyrus, rostral middle frontal gyrus, pars orbitalis, pars triangularis, pars opercularis, precentral gyrus, postcentral gyrus, supramarginal gyrus, orbital frontal gyrus, and the anterior cingulate. Visual contacts located within these regions were categorized as the fronto-parietal visual contacts (n = 63, depicted in yellow in Figure 1B). Any visually responsive contacts not falling into these specified categories were termed “other visually responsive contacts” and are depicted in white in Figure 1B, while non-visual contacts are depicted in gray. Further details can be found in ^28^.

#### Visual, visual-text and abstract-text representations using DNNs

We used three different representations that are based on different inputs:

1. Visual (images) - the visual representation of the image stimuli based on a DCNN pre-trained on ImageNet.
2. Visual descriptions (text) - visual descriptions of the image stimuli. The visual descriptions were generated by Chat-GPT^46^: We uploaded each image to chat GPT and used the following prompt: “Please describe the image based on its visual features”. In case the text included the names or the location of the image, we removed it from the text (or changed it to he/she/it). In addition, we removed text that described the image overall atmosphere. A total of 4 such sentences were removed from the descriptions of all 28 images. For visual descriptions of the images see Fig 2B (for an example) and Supplementary material (for all visual descriptions).
3. Abstract descriptions (text) – the first paragraph of the Wikipedia descriptions of the identities\places depicted in each image (see Fig 2B for an example and Supplementary material for all abstract descriptions).

We extracted the representation of each input type using the following models:

The representations of the images were extracted using a visual deep learning algorithm - VGG-16 DCNN^29^ pre-trained on ImageNet^47^, which includes 300000 images from 1000 object categories. VGG is a commonly used network that was shown to be correlated with behavioral and neural responses to the same images^48–51^. We then extracted the embeddings of each image based on the feature vector representation in the penultimate-i.e. the last hidden layer (fc7) just before the output layer of the network, which is the representation that is used for the classification. The similarity between each pair of images was computed based on the cosine distance between these feature vectors.

The representations of the text input were extracted using a large language model - GPT2: A transformer-based language algorithm^30^. GPT is trained on a vast corpus of text to carry out various language tasks (e.g. semantic similarity, questions answering, grammatical correction etc.), often involving longer text captions. We retrieved the embeddings based on the model output layer of the last word in each description. Subsequently, we computed the similarity between each stimuli pair based on the cosine distance between the representations of these textual descriptions. We extracted GPT representation once based on the visual textual descriptions and once based on the abstract textual descriptions, resulting in two different RDMs that were generated with GPT.

#### Behavioral studies

##### Human similarity ratings – faces and places

###### Stimuli

The same stimuli that were used for the iEEG recording.

###### Participants

A total of 40 participants were recruited for this study from the Prolific platform. Twenty participants were assigned to each of the two experimental conditions: Visual similarity based on images and semantic similarity based on names (mean age 29 (sd = 2.9), 20 females, 1 preferred not to say). The participants were paid 6 GBP for their participation in the experiment (9 GBP/hour). They gave informed consent prior to the experiment. The study was approved by the ethics committee of Tel Aviv University. We chose this sample size based on previous similarity tasks (Shoham et al)

###### Procedure

Participants rated the visual or semantic similarity of all possible pairs of the 28 items (378 pairs).

*Visual similarity rating:* Each trial presented one pair of images of two different items, and the participants were asked to rate the visual similarity between the items. The pairs were presented on the screen one at a time, above a similarity scale (1 (very similar) - 6 (very dissimilar)) until response. Above the images, the participants saw the question “How visually similar are these images?”. The participants selected the similarity score with the mouse. The next pair was presented 1 second after their response. The participants had a forced break for a minimum of 10 seconds after 80 pairs were presented in a row (a total of 4 breaks during the experiment). After they rated all 378 pairs, the participants were asked to indicate for each image whether they were familiar with it before the experiment. The experiment lasted about 40 minutes.

*Semantic similarity rating:* Participants were asked to rate the similarity between the names of the stimuli based on the conceptual/semantic knowledge they had about them. The timing and procedure were identical to the visual similarity task, beside that in the semantic rating task the instruction was only presented at the beginning and there was no line of instruction in each trial.

###### Trials’ exclusion

We excluded trials if the participant was unfamiliar with one of the items. Participants who were unfamiliar with 30% (or more) of the identities were excluded from the analysis. We excluded trials shorter than 200 ms and longer than 30,000 ms (30 seconds) based on the assumption that participants did not perform the task well on such trials. In the visual similarity tasks, 23% of the trials were excluded (33 trials due to RT longer than 30 seconds and the rest based on the familiarity criterion). In the semantic similarity rating task, 15% trials were excluded (50 trials due to RT longer than 30 seconds and the rest based on the familiarity criterion).

##### Text image matching task – faces and places

###### Stimuli

The same stimuli that were used for the iEEG recording.

###### Participants

A total of 40 participants were recruited for this study from the Prolific platform. Twenty participants were assigned to each of the two conditions: visual text matching (mean age 33 (sd = 3.4), 5 females) or abstract text matching (mean age 32 (sd = 3.4), 10 females). The participants were paid 2.25 GBP for their participation in the experiment (9 GBP/hour). They gave informed consent prior to the experiment. The study was approved by the ethics committee of Tel Aviv University

###### Procedure

Participants matched each visual or abstract description of all 28 images to one image out of 14 images (the images from the same category, faces or places). They were presented with one text at a time and were asked to choose the image that best matches each text. The images were presented simultaneously, and the participants selected the number of images that best matched the text. After making their choice they were presented with the next text trial.

At the end of the study, they were asked to indicate whether they were familiar with the faces/places before the study. The experiment lasted about 15 minutes.

We did not exclude trial based on any criteria.

#### Human similarity ratings – scenes

15 color images of indoor and outdoor scenes (see Supplementary Materials for all scene images and their textual descriptions).

#### Visual, visual-text and abstract-text representations using DNNs

We used three different representations that are based on different inputs:

1. Visual (images) - the visual representation of the image stimuli based on a DCNN pre-trained on ImageNet.
2. Visual descriptions (text) - Similarly to set 1, the visual descriptions were generated by Chat-GPT^46^. In case the text included the name of the scene, or the words “the image shows” we removed it from the text. For visual descriptions of the images see Fig 6A (for an example) and Supplementary material (for all visual descriptions).
3. Abstract descriptions (text) – the first paragraph of the Wikipedia descriptions of the scene depicted in each image (see Fig 6A for an example and Supplementary material for all abstract descriptions).

We extracted the representation of each input type using the following models:

The representations of the images were extracted using VGG-16 DCNN^29^ pre-trained on ImageNet^47^ (similar to the familiar face and place stimuli).

The representations of the text input were extracted using a large language model – SGPT^52^: GPT Sentence Embeddings for Semantic Search is a recent natural language processing (NLP) algorithm that is first pre-trained to predict the next word in a sentence similar to other NLP algorithms and use contrastive fine-tuning to create similar representations for pairs of sentences that describe the same content^52^. This model was chosen because the correlations of its representations with semantic similarity ratings (r = xxx) of the stimuli was higher than the representations generated by GPT-2 (r = xxx). We extracted the embeddings of the Wikipedia definition based on the 1.3B parameters bi-encoder’s output layer and computed the similarity between each scenes pair based on the cosine distance between these representations.

#### Behavioral data: Human similarity ratings

##### Participants

A total of 40 participants were recruited for this study from the Prolific platform. Twenty participants were assigned to each of the two experimental conditions: Visual similarity based on images and semantic similarity based on names (mean age 30 (sd = 2.9), 18 females). The participants were paid 6 GBP for their participation in the experiment (9 GBP/hour). They gave informed consent prior to the experiment. The study was approved by the ethics committee of Tel Aviv University. We chose this sample size based on previous similarity tasks (Shoham et al)

##### Procedure

Participants rated the visual or semantic similarity of all possible pairs of the 15 items (105 pairs). The timing and procedure of both tasks was identical to the behavioral study described for the familiar faces and places described above except that the instructions were presented at the beginning and with no reminder on each trial.

##### Trials’ exclusion

We excluded trials shorter than 200 ms and longer than 30,000 ms (30 seconds) based on the assumption that participants did not perform the task well on such trials. In the visual similarity tasks, 0.4% of the trials were excluded. In the semantic similarity rating task, 0.4% of the trials were excluded.

#### Data Analysis

We used Representational Similarity Analysis (RSA) to predict the neural and behavioral measures based on the DNNs representations. We created a dissimilarity matrix (RDM) of all possible stimuli pairs within the pool of the 28 stimuli (a total of 378 pairs), separately for high-level visual cortex and the fronto-parietal ROIs, for the visual, visual-text and abstract-text representations and for each participant in the visual and semantic behavioral ratings tasks. We performed the same analyses for the scene images (Figure 6). (15 Images, 105 pairs).

##### Correlation between the DNN representations and brain ROIs

We created a representational dissimilarity matrix (RDM) based on the iEEG responses for the different stimuli. For each stimulus, we calculated the average response (mean amplitude of the broad band high frequencies of the iEEG signal in each contact over the four stimulus repetitions, within the 0.1 to 0.4 second window after stimulus onset) for each contact in the high-visual content-selective and fronto-parietal ROI^28^ (for the time analysis reported in Figure 4, we did not average across time and used each time point separately, apart from that, we calculated everything the same). We used the averaged iEEG derived population vector as the response pattern of each stimulus within each ROI. The dissimilarity between the neuronal representations of two stimuli was represented as one minus the correlation between their activation patterns. We first computed the Pearson correlation between the DNNs’ RDMs and the RDMs of the ventro-temporal and fronto-parietal visual contacts. Then, to further assess the unique contribution of each DNN-based representation to the neural responses we computed the partial correlation of each DNN while holding the other DNNs constant.

To test the significance of a correlation/partial correlation value of a single network in a specific ROI we used a permutation test, in which the similarity scores of the neural responses were shuffled and the DNNs’ similarity scores (between pair of stimuli) were held constant. For each analysis and each ROI, we shuffled the similarity between pairs of stimuli in 10000 iterations. Then we calculated the correlation/partial correlation (respectively) of the network with the patients’ RDM (based on the shuffled similarities). P-value assigned to each network was the proportion of iterations in which the original correlations/partial correlations (based on the real similarities) were greater than the value that was calculated in each iteration. P values were adjusted according to Phipson and Smyth^53^ correction for permutation tests, and then they were FDR corrected for all networks that were tested in the same analysis, for each ROI separately. Then we used a two-tailed test to infer significance. We also tested the significance of the difference between each pair of two DNN-based representations. To do so we used a permutation test in which the networks-based similarity scores were held constant, but we shuffled the similarity scores that are brain based. For each analysis and each ROI, we shuffled the similarity between pairs of stimuli in 10000 iterations. We first calculated the correlation/partial correlation (respectively) of each network with the neural RDM (based on the shuffled similarities). We then calculated the difference between each two network representations correlations/partial correlations (with the neural RDMs). P value assigned to each comparison of a pair of networks was the proportion of iterations in which the original difference (in the same direction) in correlations/partial correlation, based on the real similarities, was greater (or smaller) than the value that was calculated in each iteration. P values were adjusted according to^53^ correction for permutation tests, and then they were FDR corrected for all networks that were tested in the same analysis, for each ROI separately. Then we used a two-tailed test to infer significance. For the time analysis reported in Figure 4, we performed the permutation test for each time point separately, and then FDR corrected for all time points together.

##### Correlation between the DNNs representations and behavioral similarity ratings

We created a representational dissimilarity matrix (RDM) for each participant based on their similarity ratings. We computed the partial correlations between the RDM of each participant and the RDM of the DNNs representations. All correlations were fisher transformed. To test for significance, ANOVA, post hoc comparisons were performed on the Fisher’s z-transformed partial correlations using two-sided null-hypothesis tests. FDR was applied to correct for multiple comparisons for all post-hoc tests in each analysis separately.

## Acknowledgments

The study was funded by a CIFAR BMC fellowship to R. Malach, ISF grant no.917/21 to G. Yovel and the Sieratzki Institute for Advances in Neuroscience scholarship to A. Shoham.

## Supplementary data

**Table 1:**
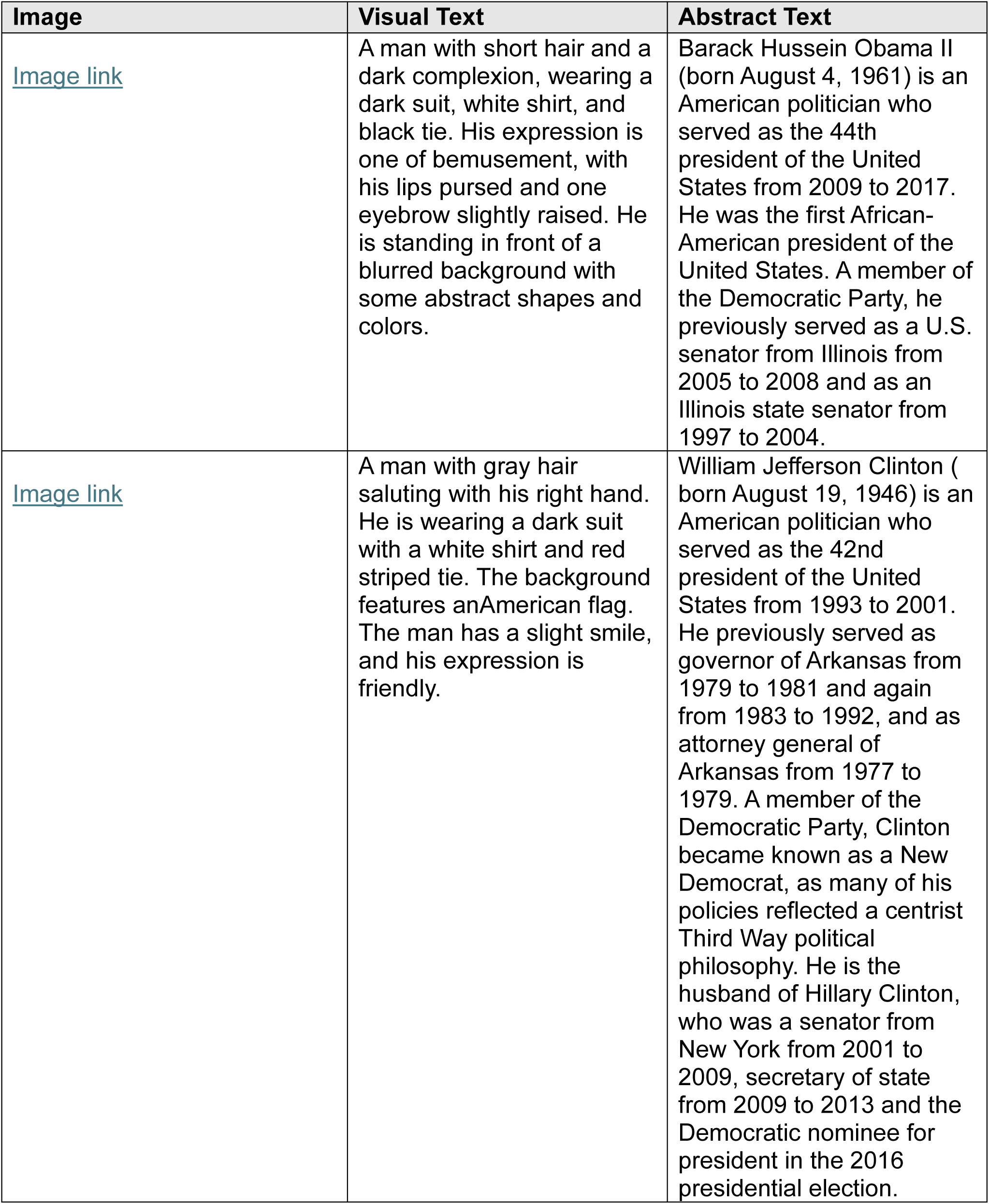
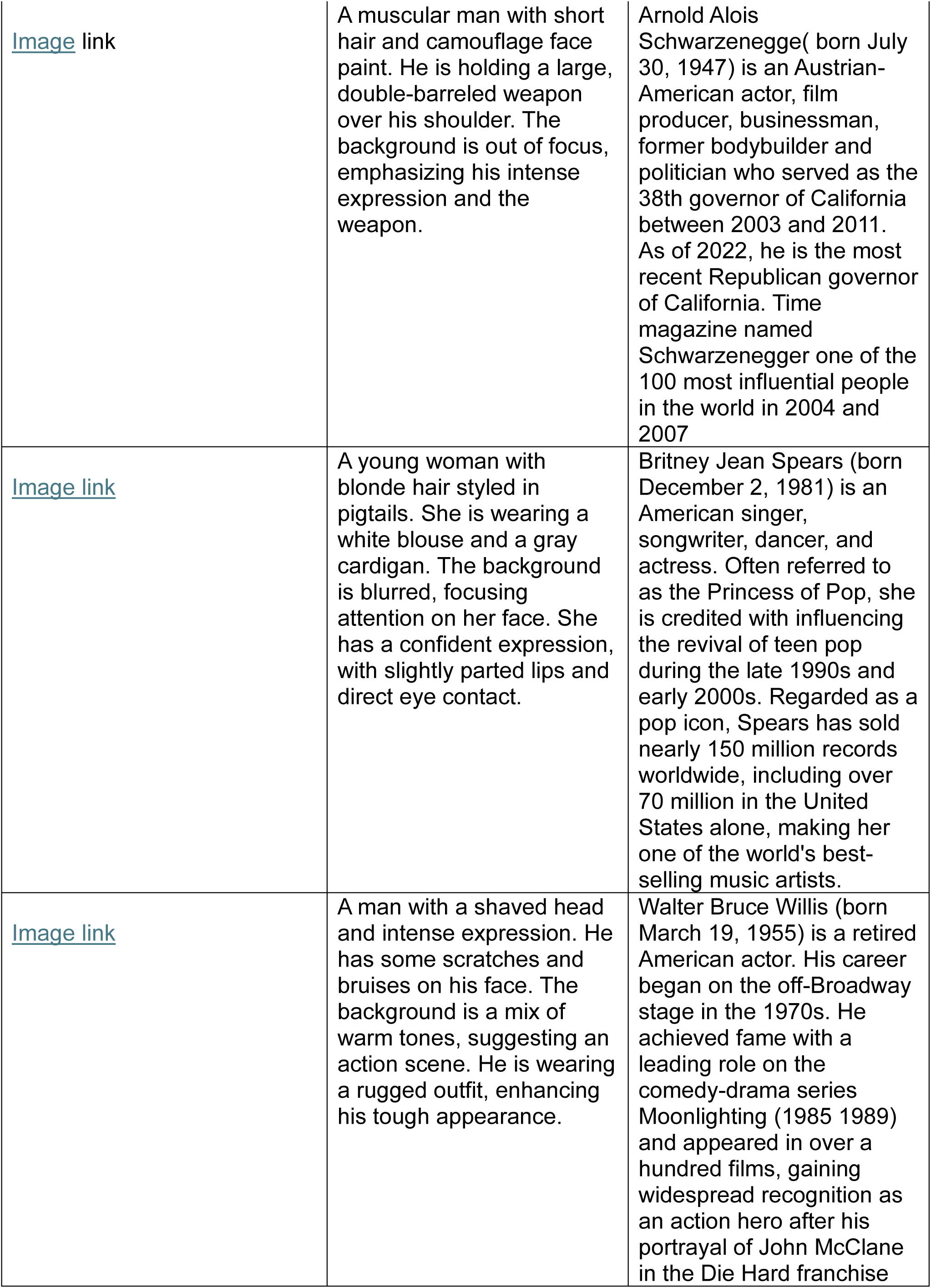
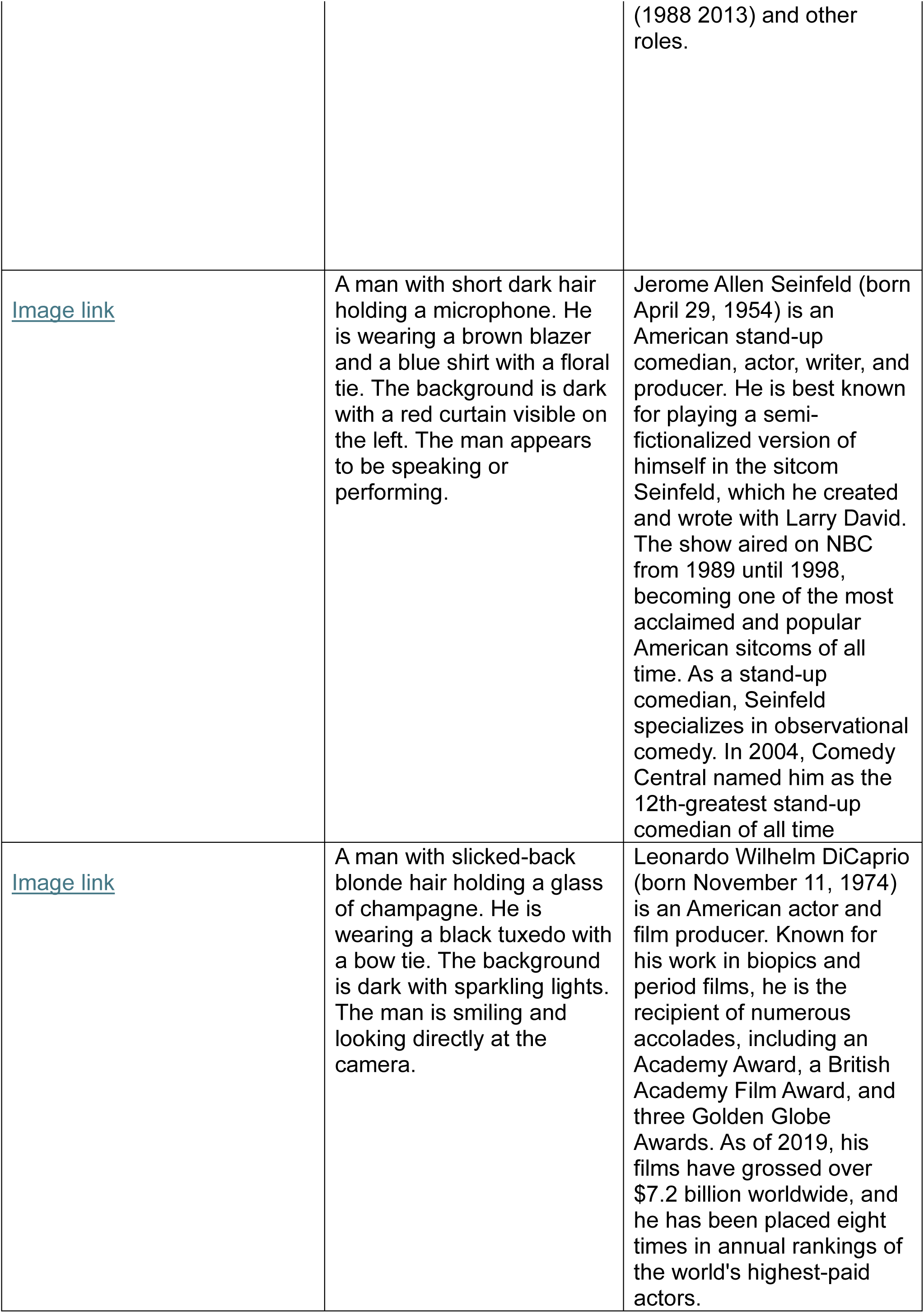
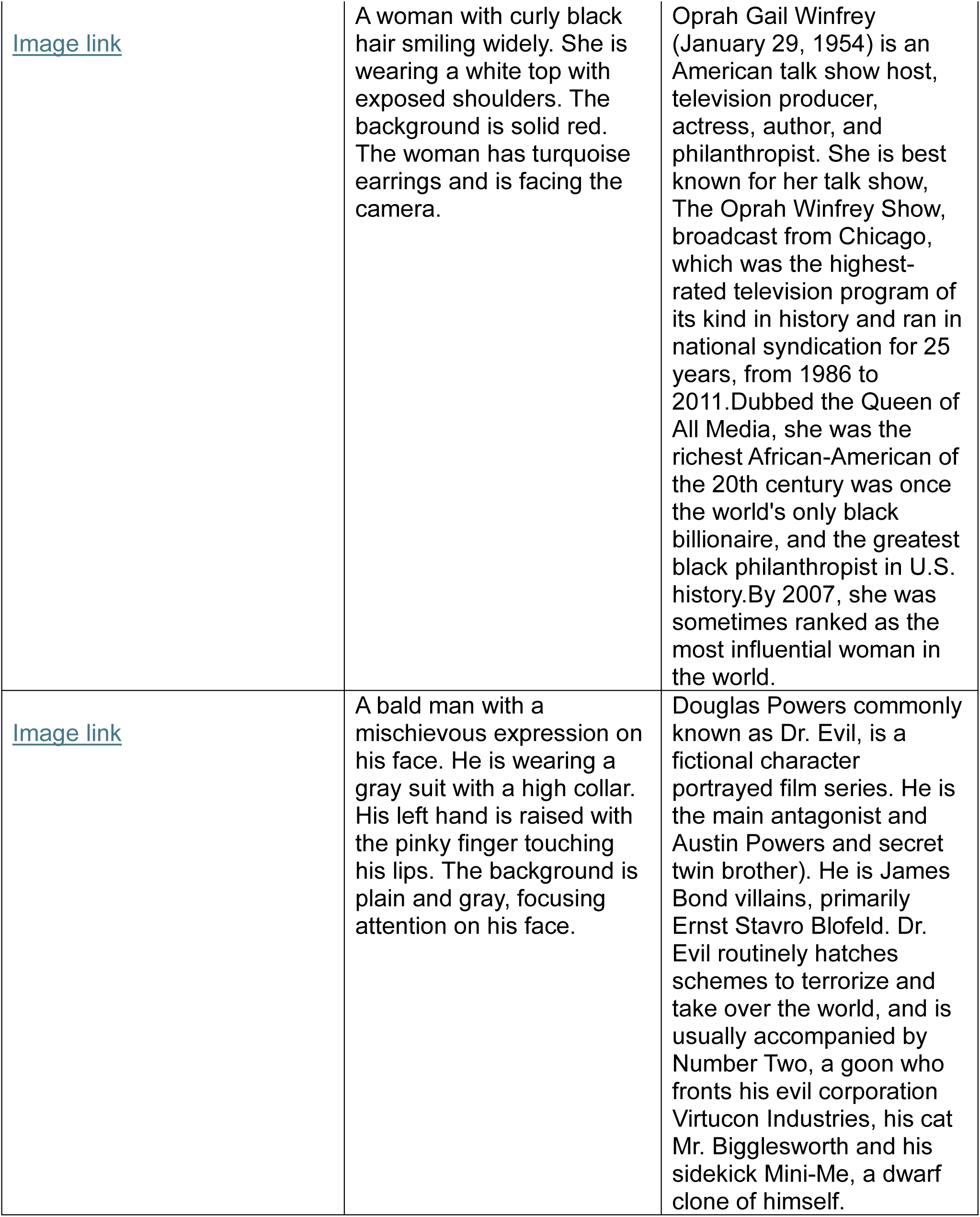
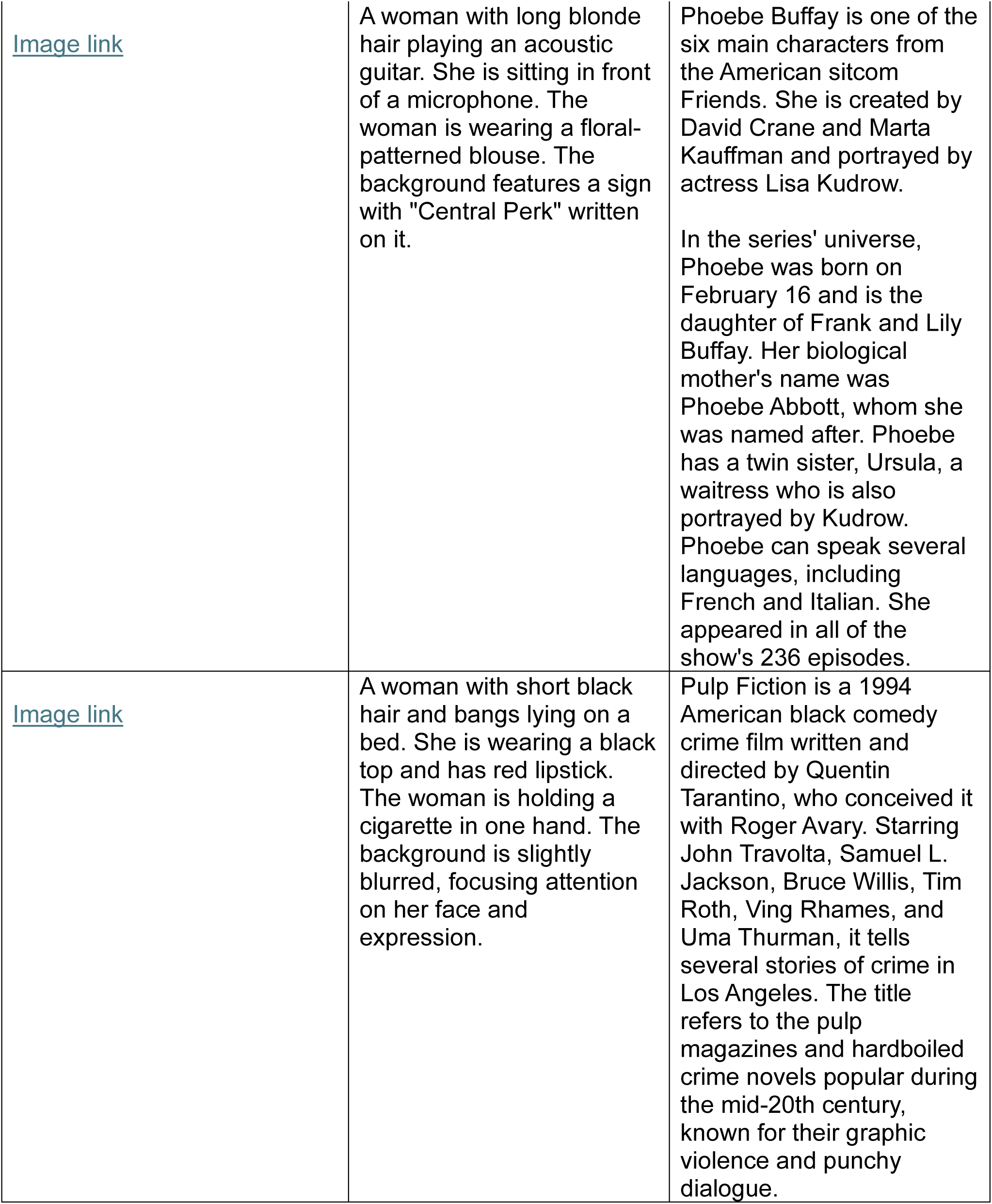
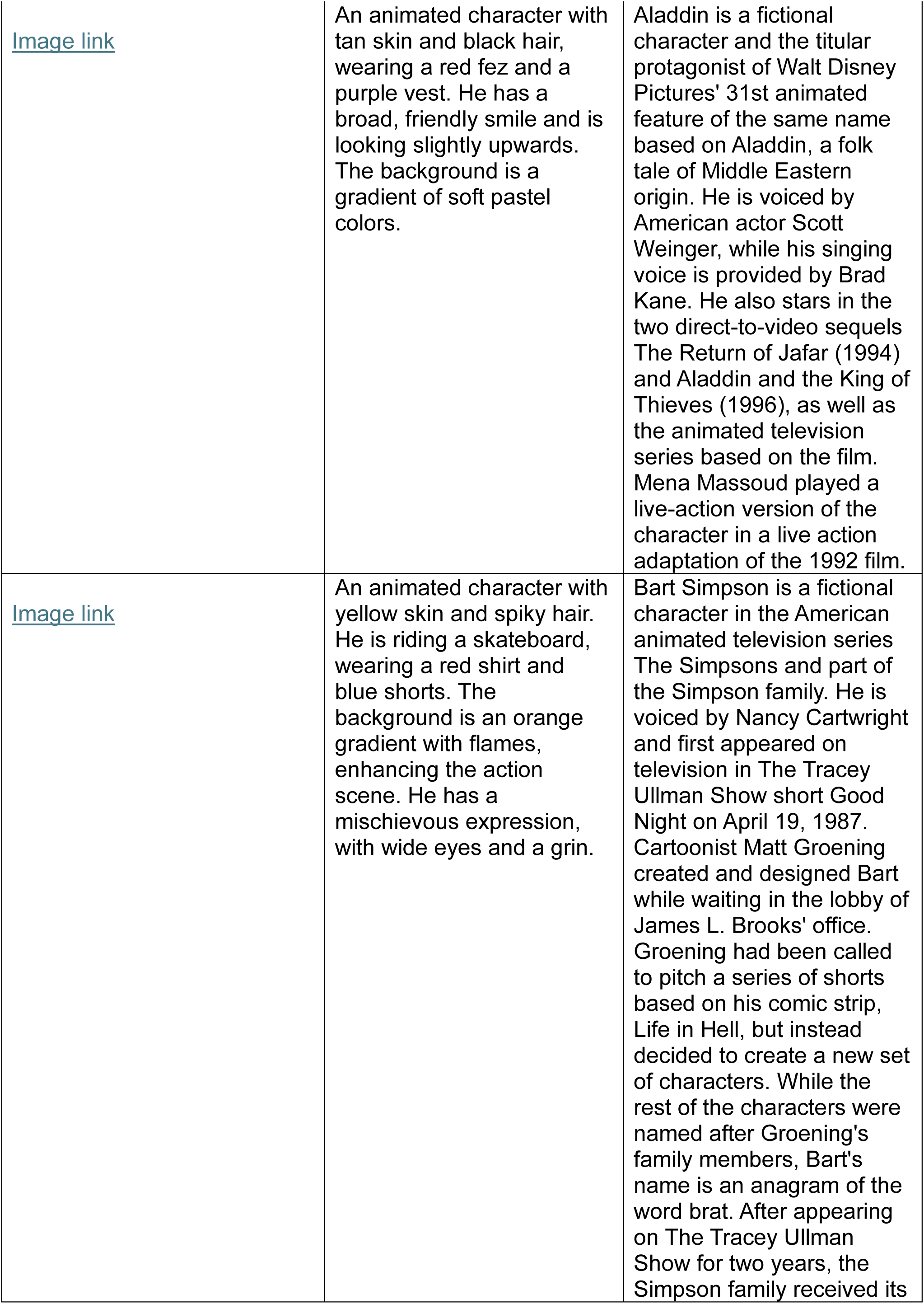
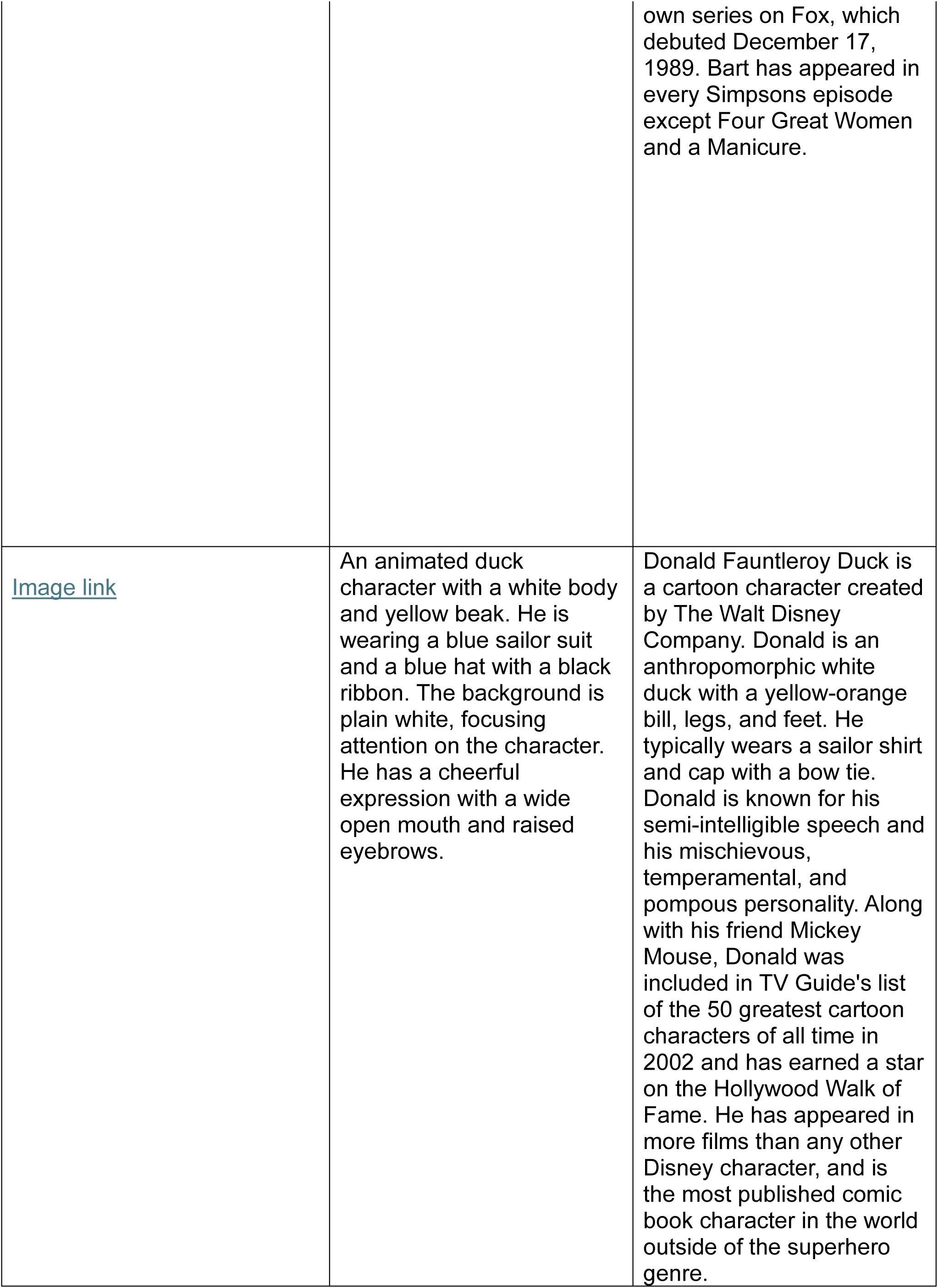
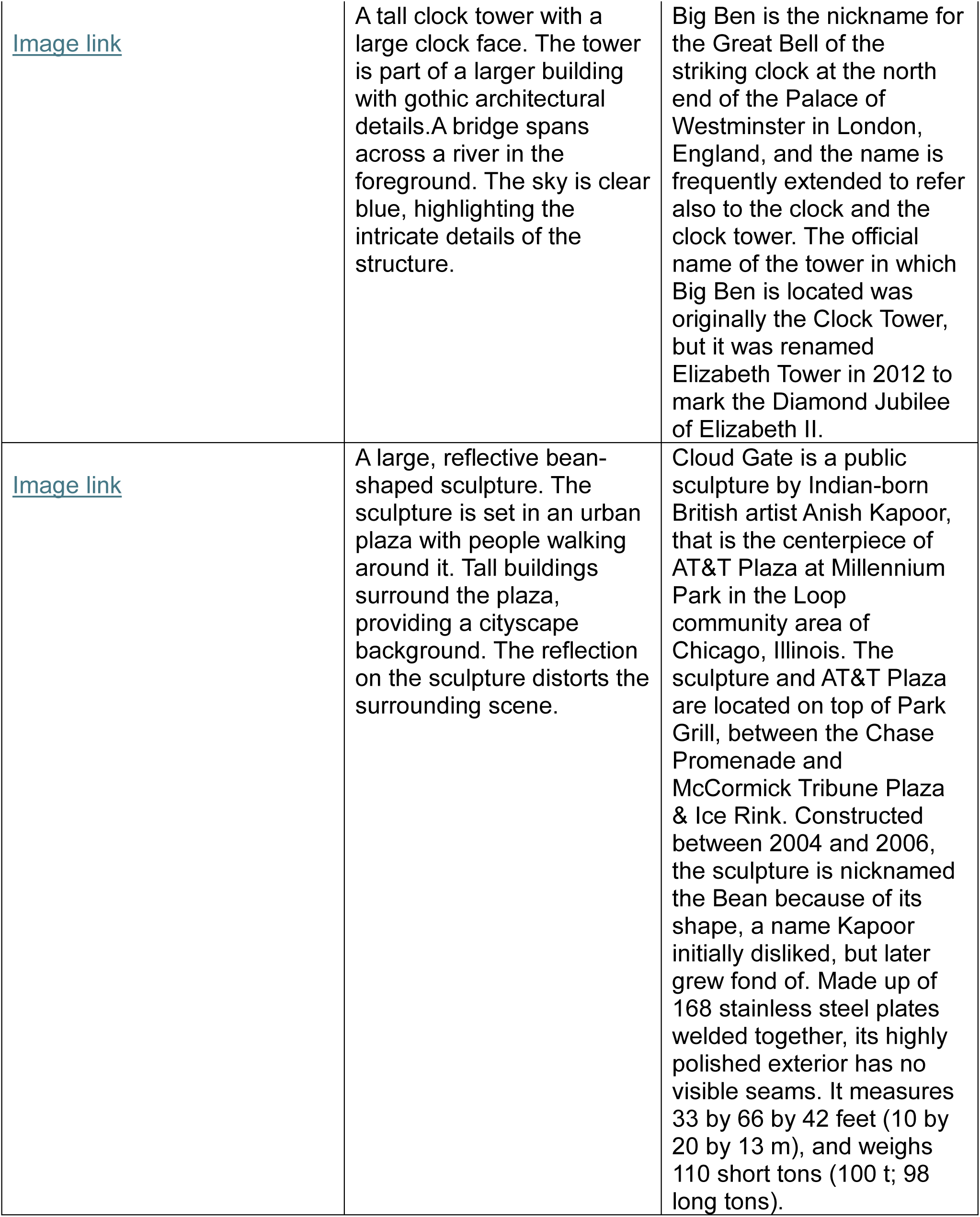
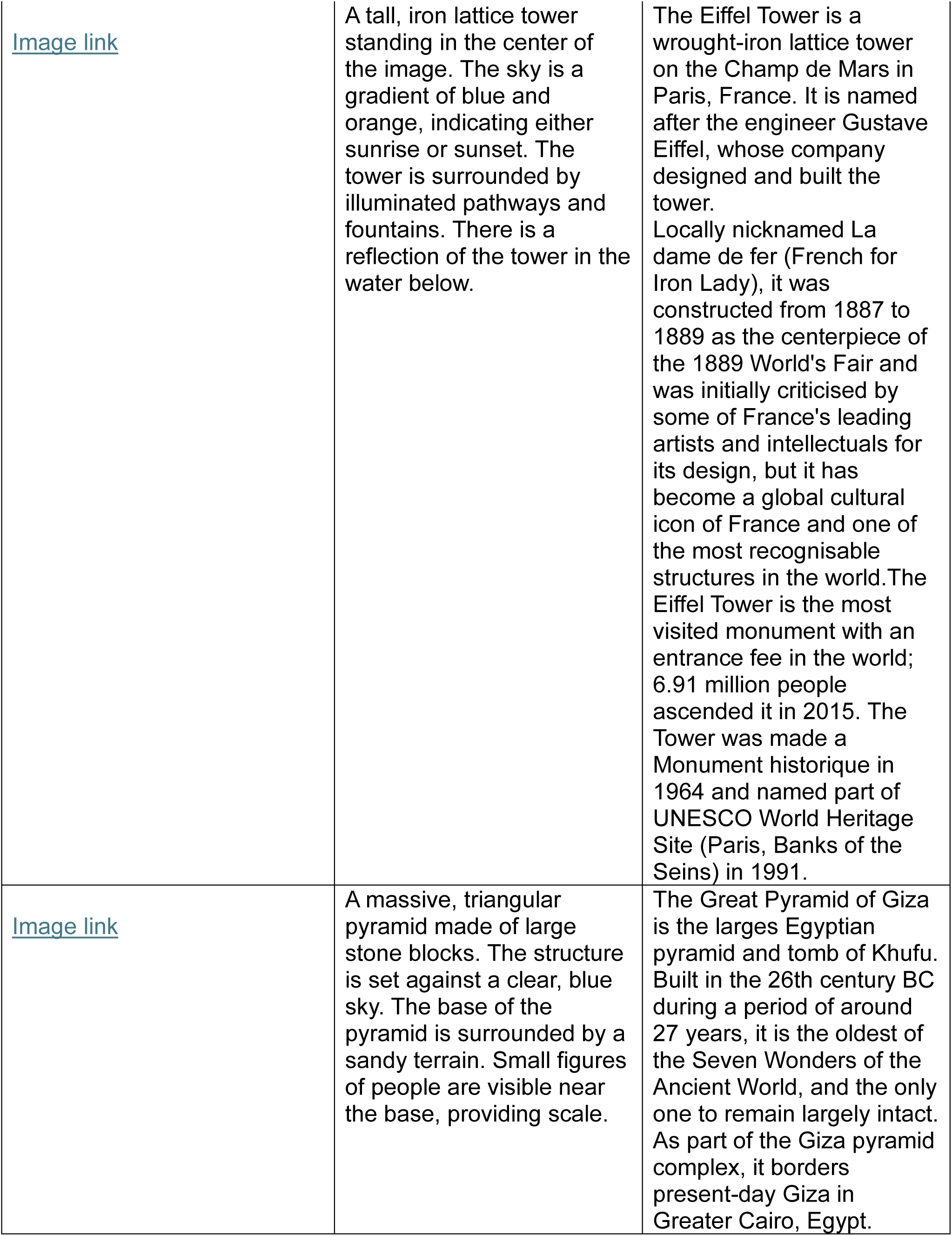
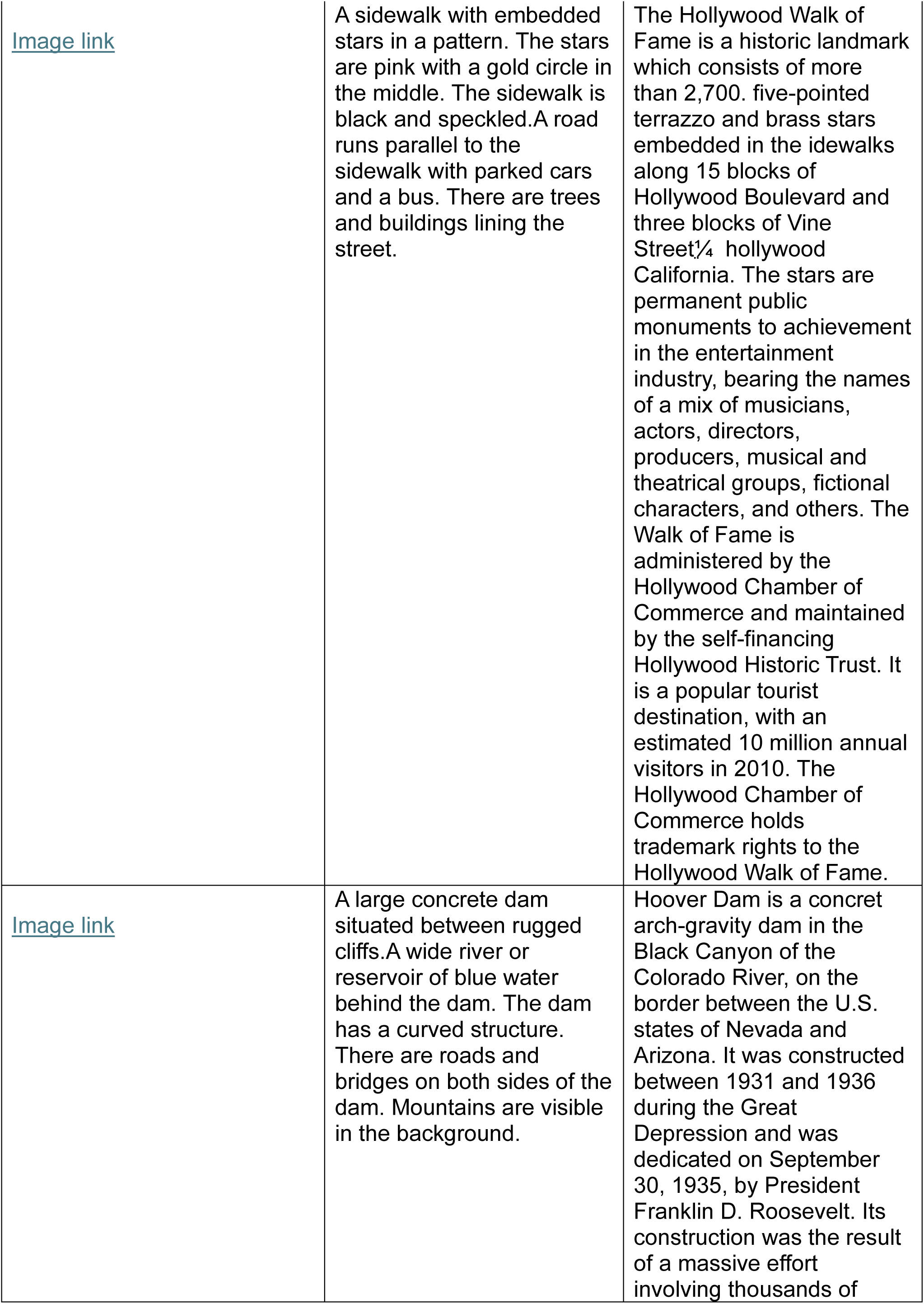
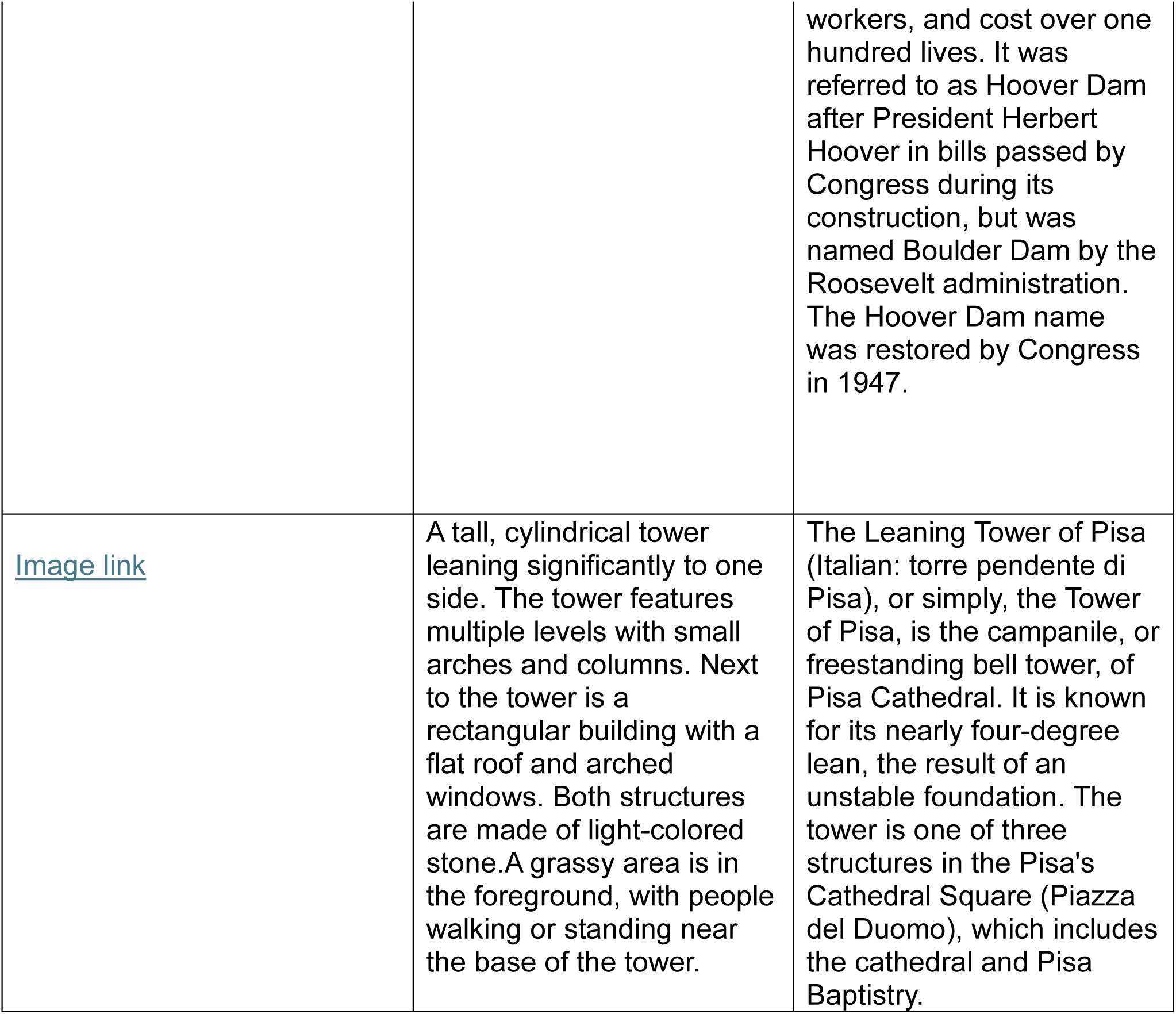
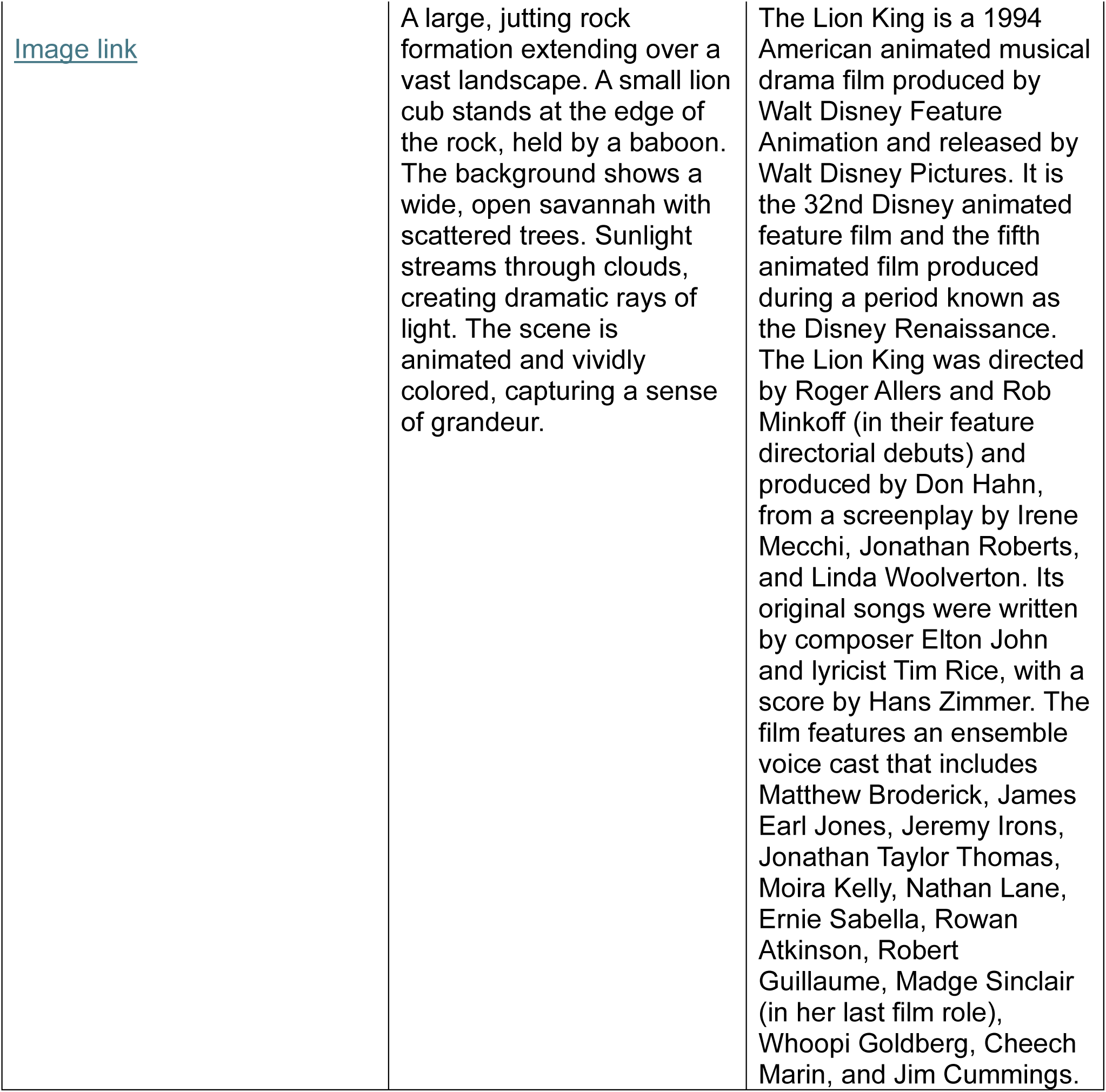
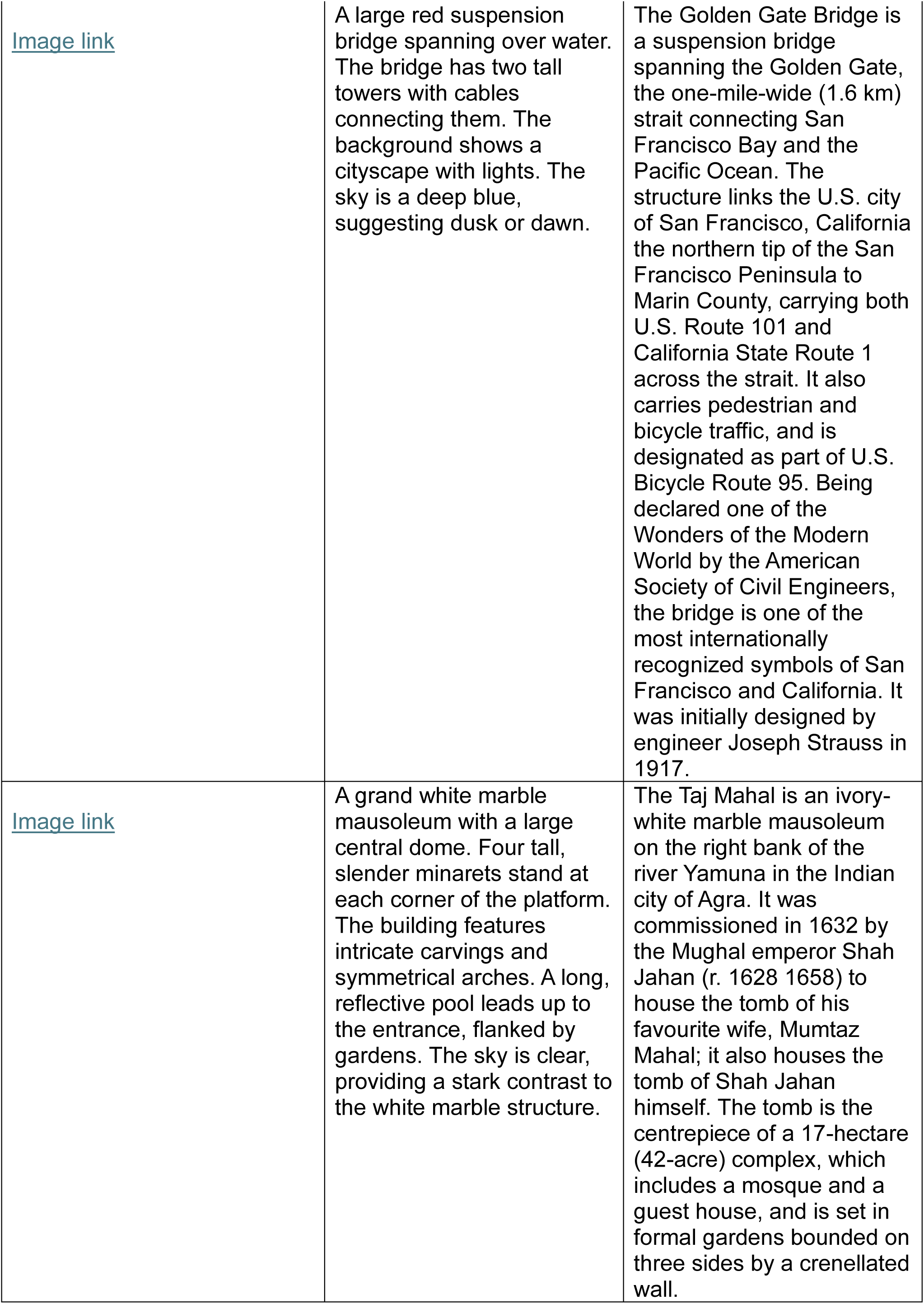
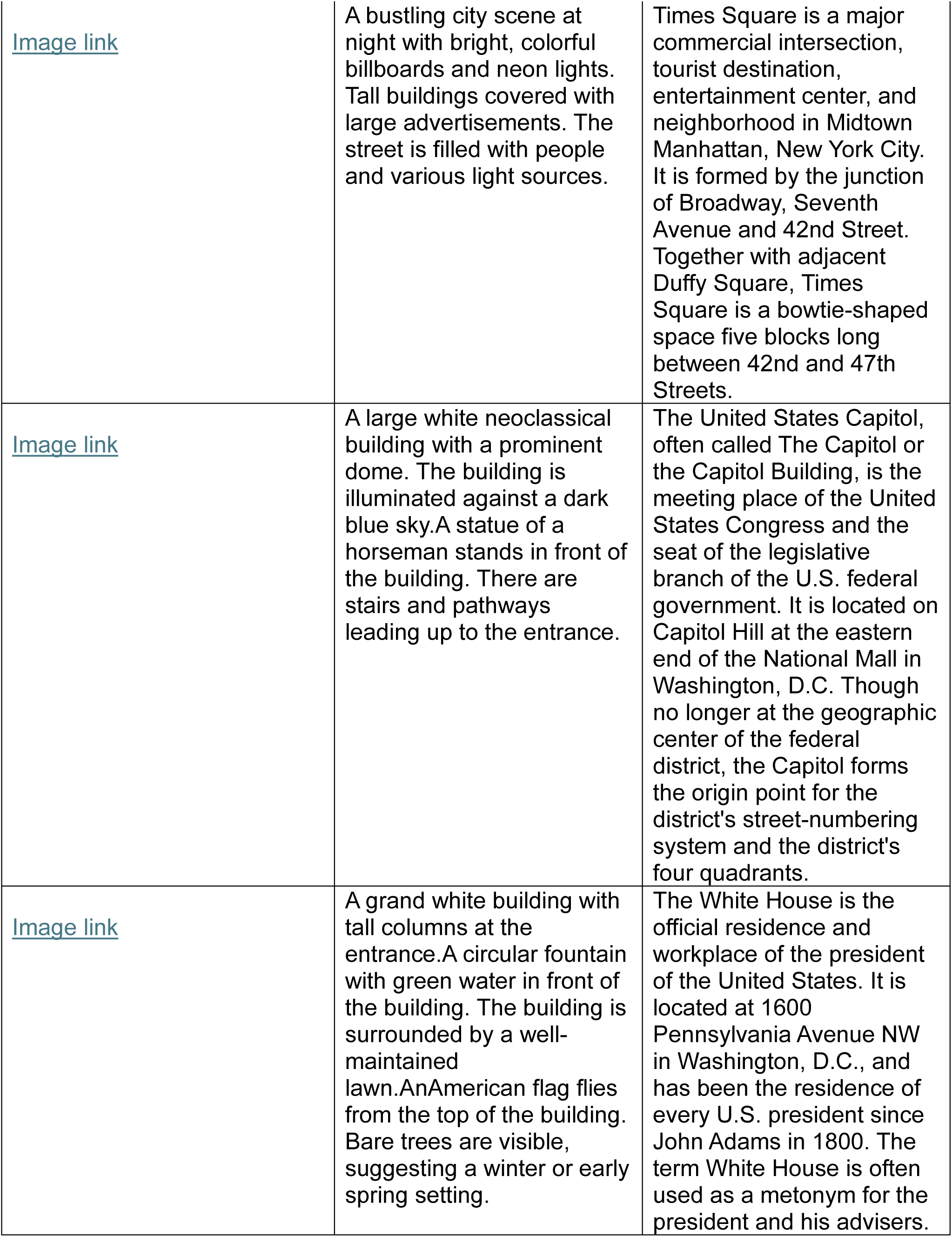
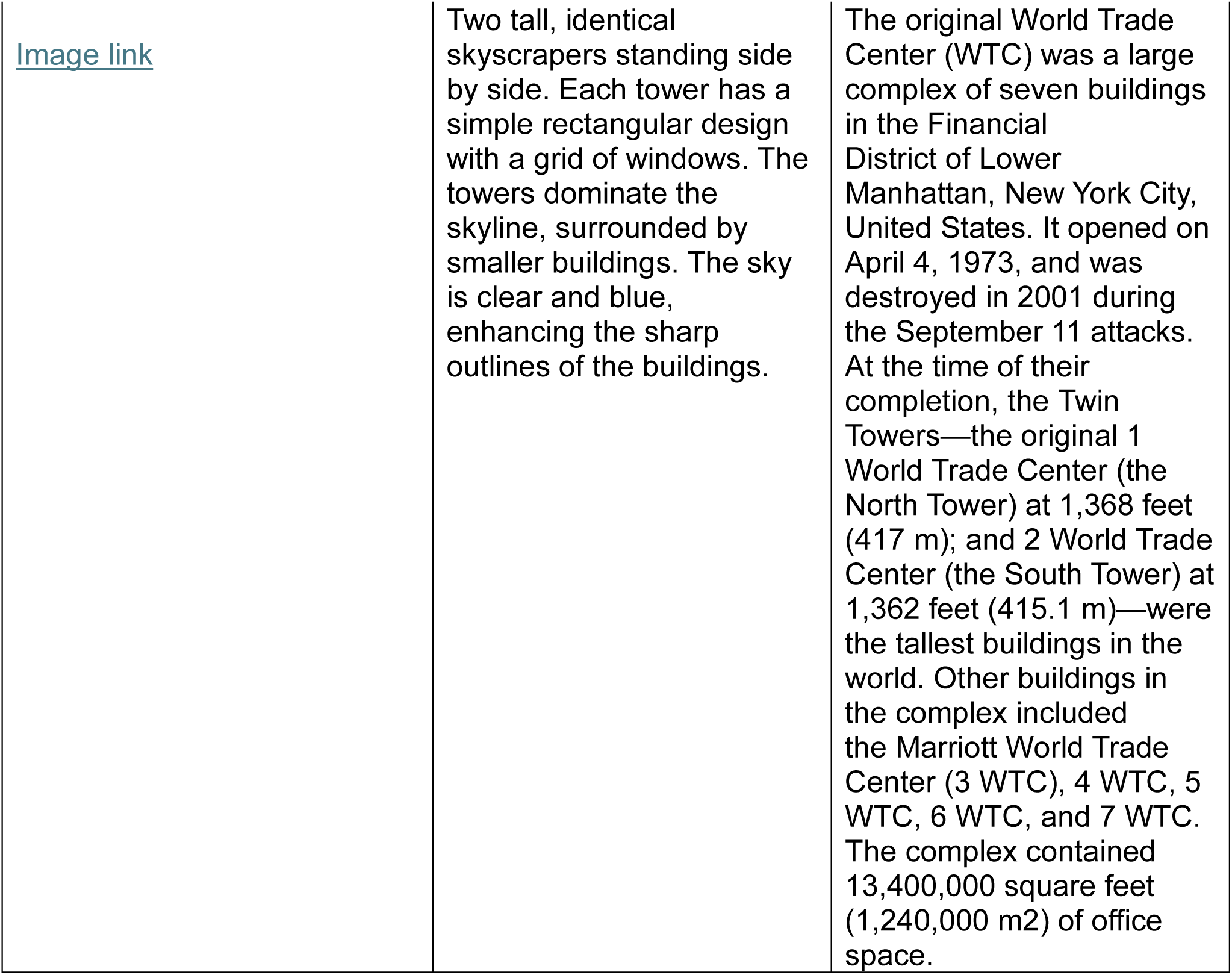
Familiar faces and places images:

**Table 2:**
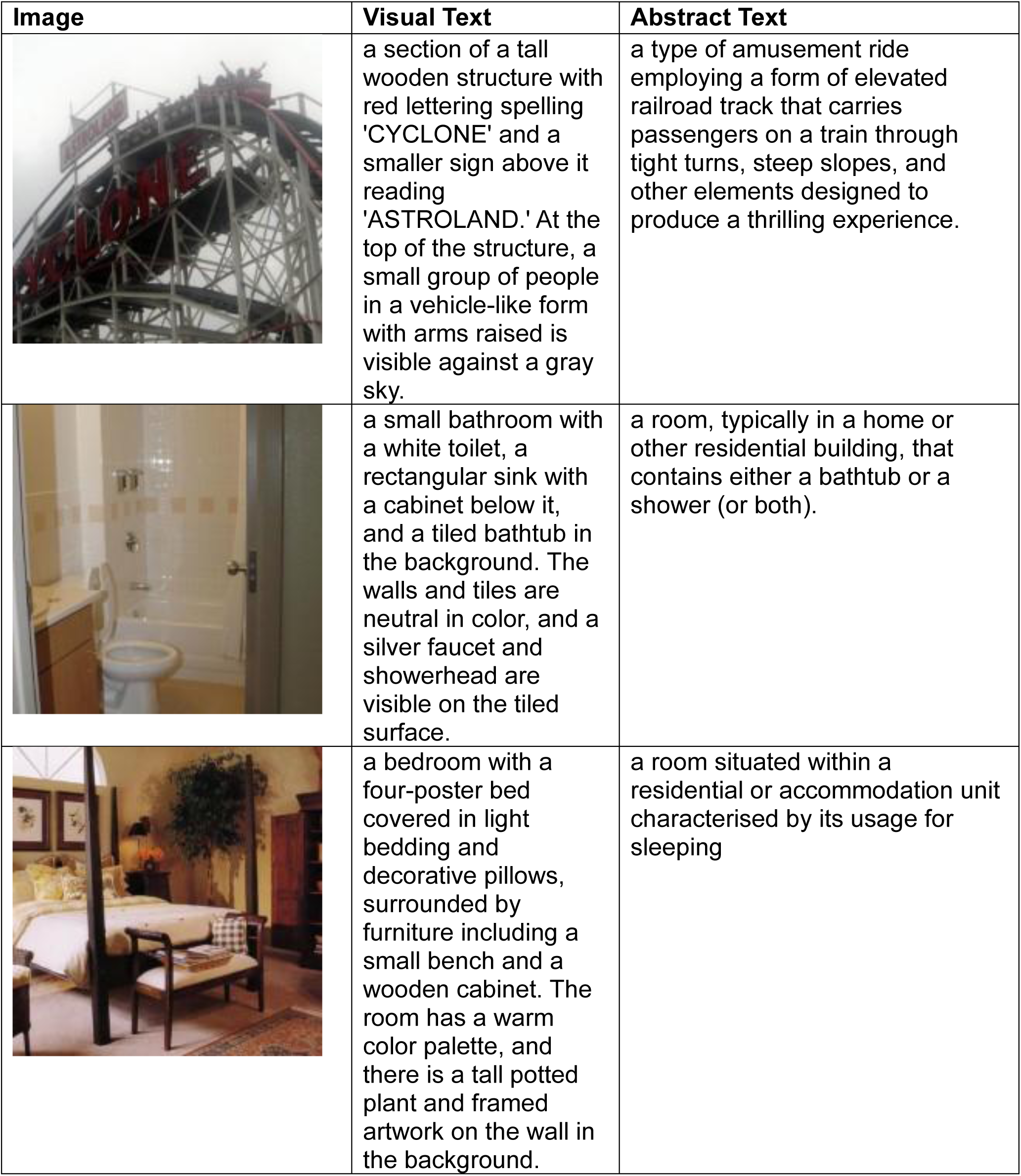
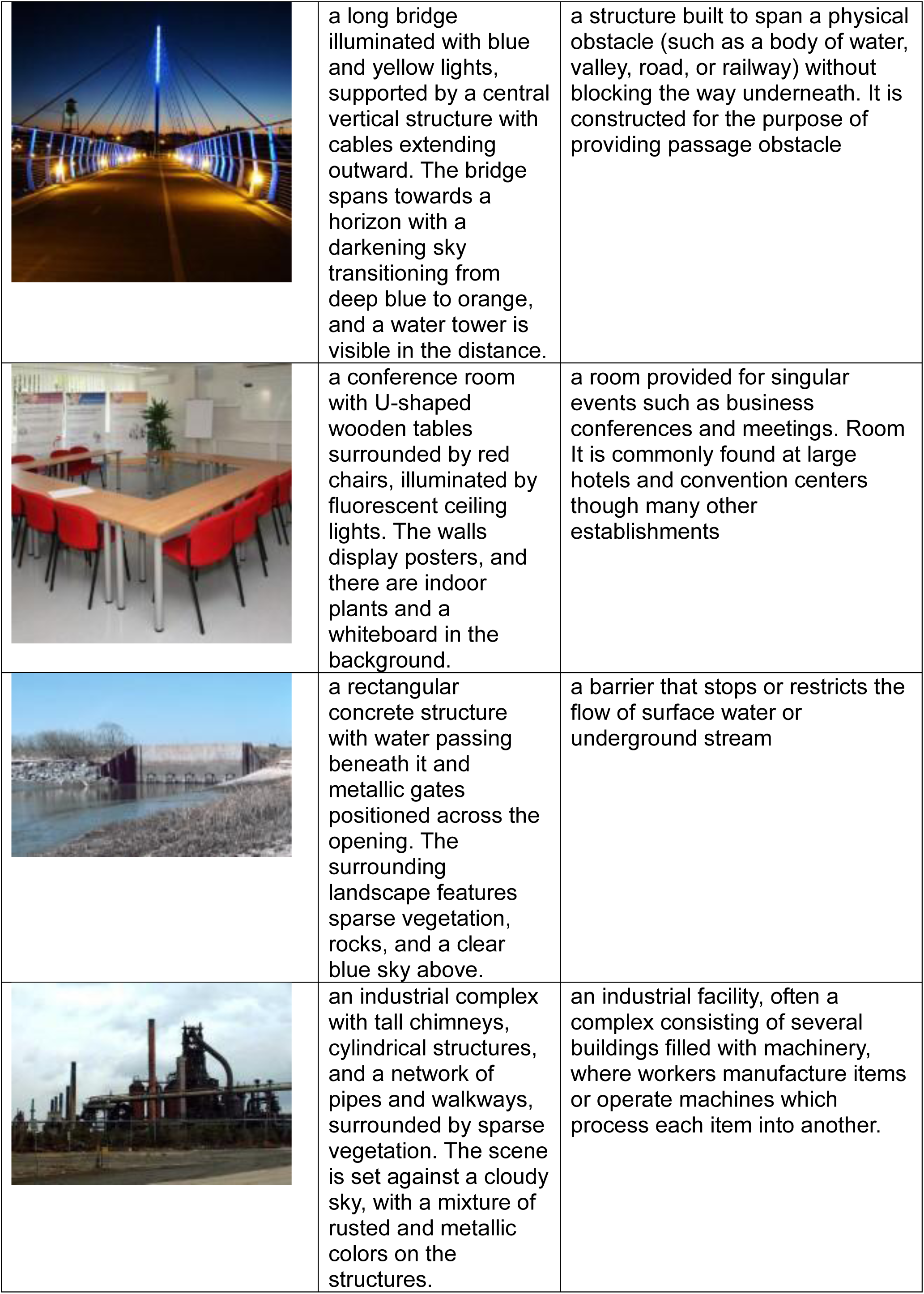
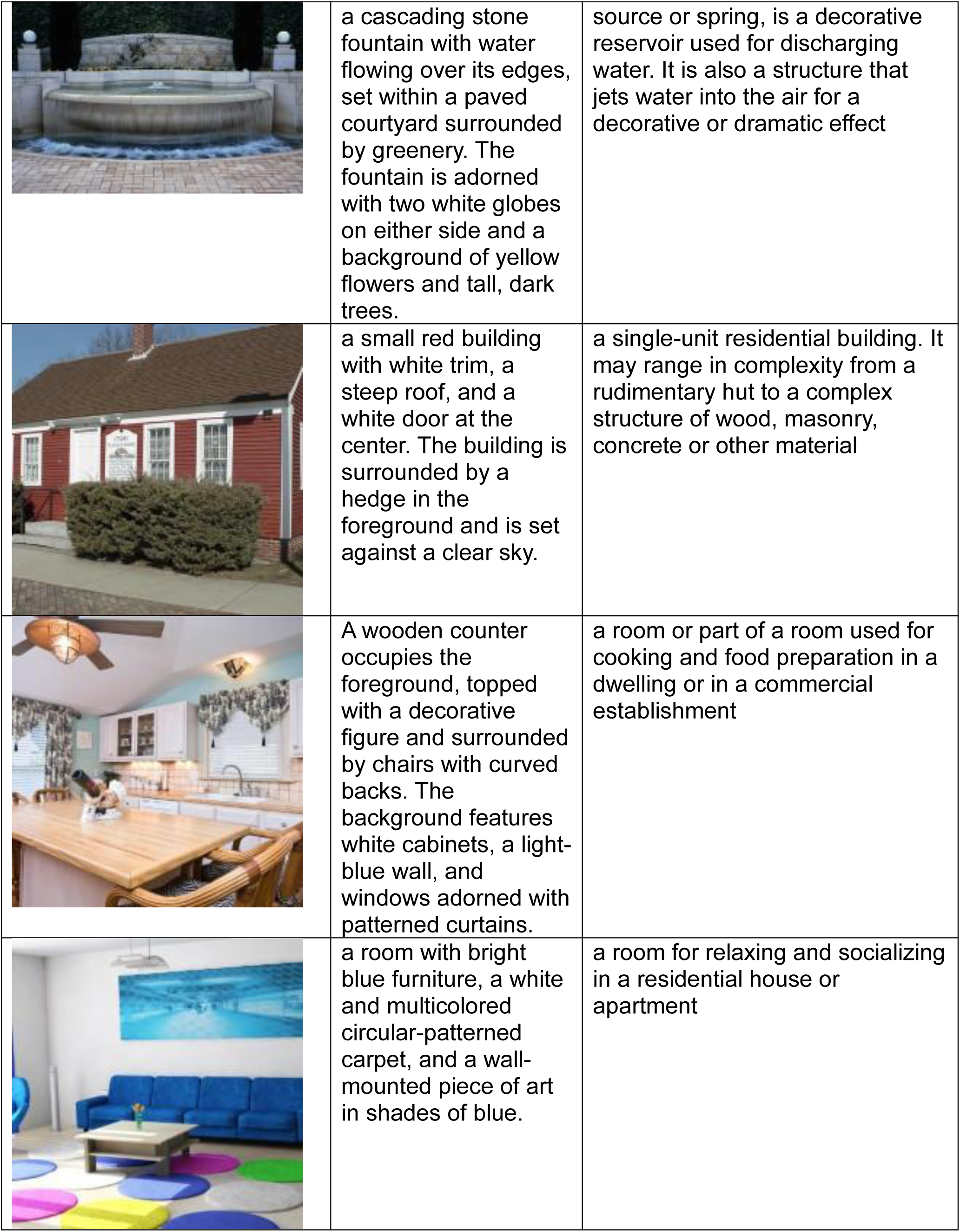
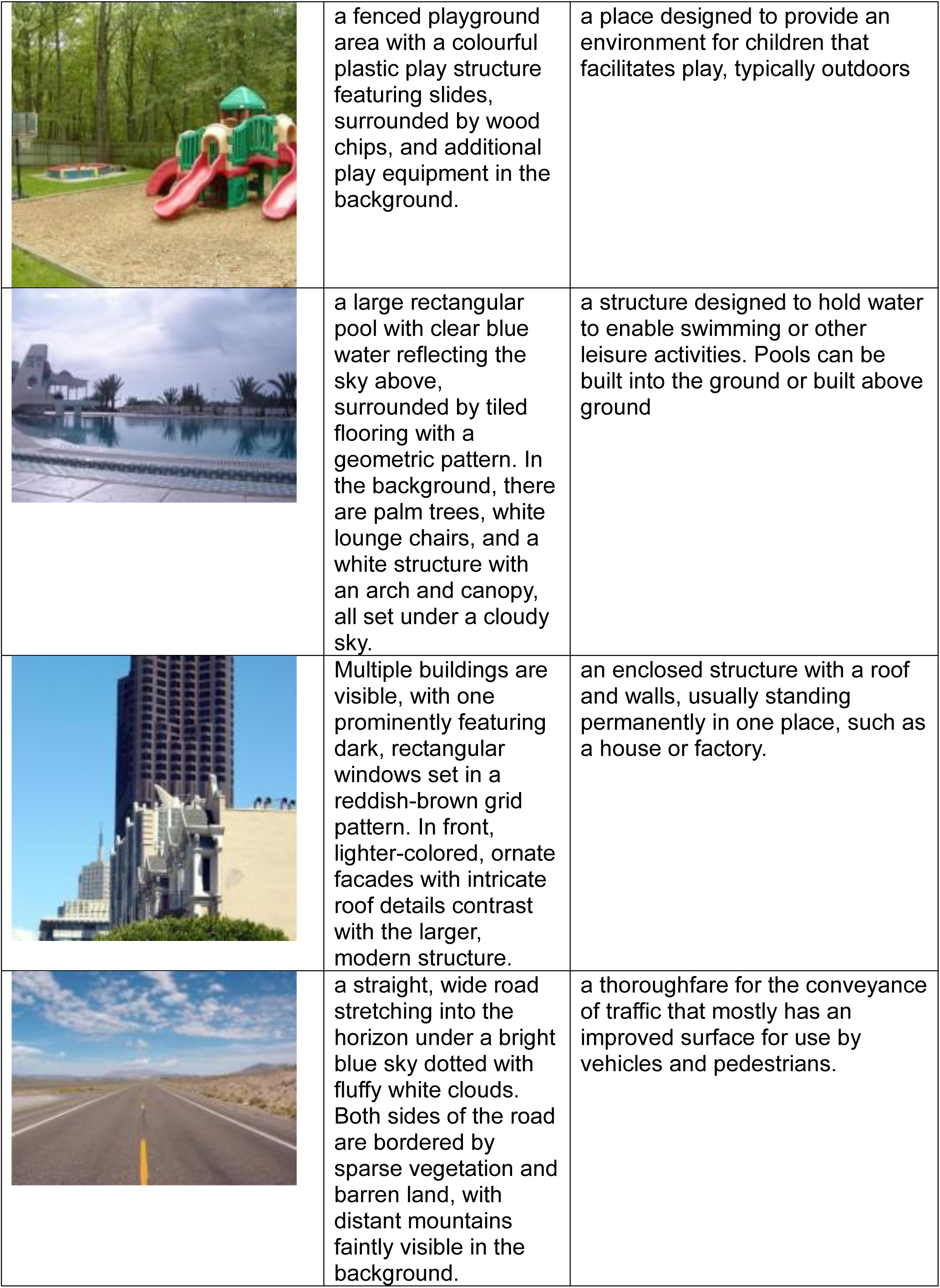
Scenes images: Taken from Bainbridge et al. ^31^.

| Image | Visual Text | Abstract Text |
| --- | --- | --- |
|  | a section of a tall wooden structure with red lettering spelling 'CYCLONE' and a smaller sign above it reading 'ASTROLAND.' At the top of the structure, a small group of people in a vehicle-like form with arms raised is visible against a gray sky. | a type of amusement ride employing a form of elevated railroad track that carries passengers on a train through tight turns, steep slopes, and other elements designed to produce a thrilling experience. |
|  | a small bathroom with a white toilet, a rectangular sink with a cabinet below it, and a tiled bathtub in the background. The walls and tiles are neutral in color, and a silver faucet and showerhead are visible on the tiled surface. | a room, typically in a home or other residential building, that contains either a bathtub or a shower (or both). |
|  | a bedroom with a four-poster bed covered in light bedding and decorative pillows, surrounded by furniture including a small bench and a wooden cabinet. The room has a warm color palette, and there is a tall potted plant and framed artwork on the wall in the background. | a room situated within a residential or accommodation unit characterised by its usage for sleeping |
|  | a long bridge illuminated with blue and yellow lights, supported by a central vertical structure with cables extending outward. The bridge spans towards a horizon with a darkening sky transitioning from deep blue to orange, and a water tower is visible in the distance. | a structure built to span a physical obstacle (such as a body of water, valley, road, or railway) without blocking the way underneath. It is constructed for the purpose of providing passage obstacle |
|  | a conference room with U-shaped wooden tables surrounded by red chairs, illuminated by fluorescent ceiling lights. The walls display posters, and there are indoor plants and a whiteboard in the background. | a room provided for singular events such as business conferences and meetings. Room It is commonly found at large hotels and convention centers though many other establishments |
|  | a rectangular concrete structure with water passing beneath it and metallic gates positioned across the opening. The surrounding landscape features sparse vegetation, rocks, and a clear blue sky above. | a barrier that stops or restricts the flow of surface water or underground stream |
|  | an industrial complex with tall chimneys, cylindrical structures, and a network of pipes and walkways, surrounded by sparse vegetation. The scene is set against a cloudy sky, with a mixture of rusted and metallic colors on the structures. | an industrial facility, often a complex consisting of several buildings filled with machinery, where workers manufacture items or operate machines which process each item into another. |
|  | a cascading stone fountain with water flowing over its edges, set within a paved courtyard surrounded by greenery. The fountain is adorned with two white globes on either side and a background of yellow flowers and tall, dark trees. | source or spring, is a decorative reservoir used for discharging water. It is also a structure that jets water into the air for a decorative or dramatic effect |
|  | a small red building with white trim, a steep roof, and a white door at the center. The building is surrounded by a hedge in the foreground and is set against a clear sky. | a single-unit residential building. It may range in complexity from a rudimentary hut to a complex structure of wood, masonry, concrete or other material |
|  | A wooden counter occupies the foreground, topped with a decorative figure and surrounded by chairs with curved backs. The background features white cabinets, a light-blue wall, and windows adorned with patterned curtains. | a room or part of a room used for cooking and food preparation in a dwelling or in a commercial establishment |
|  | a room with bright blue furniture, a white and multicolored circular-patterned carpet, and a wall-mounted piece of art in shades of blue. | a room for relaxing and socializing in a residential house or apartment |
|  | a fenced playground area with a colourful plastic play structure featuring slides, surrounded by wood chips, and additional play equipment in the background. | a place designed to provide an environment for children that facilitates play, typically outdoors |
|  | a large rectangular pool with clear blue water reflecting the sky above, surrounded by tiled flooring with a geometric pattern. In the background, there are palm trees, white lounge chairs, and a white structure with an arch and canopy, all set under a cloudy sky. | a structure designed to hold water to enable swimming or other leisure activities. Pools can be built into the ground or built above ground |
|  | Multiple buildings are visible, with one prominently featuring dark, rectangular windows set in a reddish-brown grid pattern. In front, lighter-colored, ornate facades with intricate roof details contrast with the larger, modern structure. | an enclosed structure with a roof and walls, usually standing permanently in one place, such as a house or factory. |
|  | a straight, wide road stretching into the horizon under a bright blue sky dotted with fluffy white clouds. Both sides of the road are bordered by sparse vegetation and barren land, with distant mountains faintly visible in the background. | a thoroughfare for the conveyance of traffic that mostly has an improved surface for use by vehicles and pedestrians. |

